# Identification of the potassium binding site in serotonin transporter SERT

**DOI:** 10.1101/2023.12.29.573628

**Authors:** Eva Hellsberg, Danila Boytsov, Qingyang Chen, Marco Niello, Michael Freissmuth, Gary Rudnick, Yuan-Wei Zhang, Walter Sandtner, Lucy R Forrest

**Author notes:** These authors contributed equally to this work (Eva Hellsberg, Danila Boytsov, Qingyang Chen).

## Abstract

Clearance of serotonin (5-hydroxytryptamine, 5-HT) from the synaptic cleft after neuronal signaling is mediated by serotonin transporter SERT, which couples this process to the movement of a Na^+^ ion down its chemical gradient. After release of 5-HT and Na^+^ into the cytoplasm, the transporter faces a rate-limiting challenge of resetting its conformation to be primed again for 5-HT and Na^+^ binding. Early studies of vesicles containing native SERT revealed that K^+^ gradients can provide an additional driving force, via K^+^ antiport. Moreover, under appropriate conditions, a H^+^ ion can replace K^+^. Intracellular K^+^ accelerates the resetting step. Structural studies of SERT have identified two binding sites for Na^+^ ions, but the K^+^ site remains enigmatic. Here, we show that K^+^ antiport can drive substrate accumulation into vesicles containing SERT extracted from a heterologous expression system, allowing us to study the residues responsible for K^+^ binding. To identify candidate binding residues, we examine many cation binding configurations using molecular dynamics simulations, predicting that K^+^ binds to the so-called Na2 site. Site directed mutagenesis of residues in this site can eliminate the ability of both K^+^ and H^+^ to drive 5-HT accumulation into vesicles and, in patch clamp recordings, prevent the acceleration of turnover rates and the formation of a channel-like state by K^+^ or H^+^. In conclusion, the Na2 site plays a pivotal role in orchestrating the sequential binding of Na^+^ and then K^+^ (or H^+^) ions to facilitate 5-HT uptake in SERT.

**Significance statement:** Neuronal signaling depends on efficient clearance of the neurotransmitter from the synaptic cleft. To this end, proteins such as serotonin transporter (SERT) leverage the gradients of Na^+^ and K^+^ ions across the cell membrane, generated by Na^+^/K^+^-ATPase. While the role of Na^+^ in neurotransmitter transport is well understood, our understanding of the role of potassium in SERT has been limited. In this study, the authors use a combination of biochemical, electrophysiological, and computational tools, to identify the Na2 site as the binding site for K^+^, shedding light on a critical aspect of neurotransmitter transport.

## Introduction

Serotonin transporter (SERT, SLC6A4) plays a pivotal role in the nervous system by regulating the synaptic levels of the neurotransmitter serotonin (5-hydroxytryptamine or 5-HT). SERT is essential for mood regulation and is, consequently, a prominent target for pharmaceutical agents, including antidepressants and drugs of abuse. 5- HT uptake across the plasma membrane is driven by a gradient of Na^+^ ions at a 1:1 stoichiometry, and requires extracellular Cl^−^ ions (1–4), similar to other solute carrier 6 (SLC6) transporters (5).

The 5-HT transport cycle is initiated by binding of Na^+^ to a conformation of SERT in which the 5-HT binding site is exposed only to the cell exterior (6) precluding Na^+^ release into the cytoplasm. Subsequent binding of extracellular Cl^−^ and 5-HT to this resting state facilitates a protein conformational change, closing the extracellular pathway and exposing the central (or S1) binding site to the cytoplasm (6–8), according to the alternating-access mechanism (9). This conformational change, along with subsequent dissociation of Na^+^ into the cytoplasm, has been detected as a rapid inward peak current in SERT-expressing cells under whole cell patch clamp (10, 11) (see Fig. 5K). The rate-determining conformational change to return to an outward-facing state (11, 12) can be accelerated by intracellular K^+^, increasing the turnover rate (12, 13).

Even when accelerated by K^+^, the return step is almost two orders of magnitude slower than the 5-HT-dependent forward step (14). Consequently, most of the transporters at the cell surface are in an inward-facing conformation during steady-state transport. In the presence of cytoplasmic K^+^, this conformation of SERT can become a conducting state whose steady, channel-like current is uncoupled from the transport cycle (11, 14). Similar to currents observed in the closely related dopamine transporter, DAT (15), this current reflects inward flux of cations, presumably Na^+^.

An interaction with intracellular K^+^ has also been reported for DAT (15, 16), though it has been more difficult to demonstrate that the K^+^ efflux is coupled directly to dopamine uptake (17).

Early studies of SERT in platelet plasma membrane vesicles (PPMV) demonstrated that an outward-directed gradient of K^+^ could drive accumulation of 5-HT with a 1:1 stoichiometry (3), consistent with its impact on rates measured by patch clamp experiments (15). The overall transport cycle then involves one Na^+^ entering the cell and one K^+^ ion exiting (see Fig. 2B), unlike Cl^−^ which, although required, apparently does not cross the membrane during the transport cycle (11). Notably, K^+^ stimulation is not required for 5-HT uptake (12), seemingly because intracellular H^+^ can substitute for K^+^ ions under appropriate conditions (18). Specifically, a transmembrane pH difference (ΔpH, more acidic inside) stimulated transport into PPMV even in the absence of a Na^+^ gradient. In addition, elevated intracellular H^+^ doubled the uptake rate, restored the substrate-induced steady current, and reduced the net positive charge appearing to cross the membrane (11), mimicking the phenotype of intracellular K^+^. In essence, K^+^ or H^+^ compete with Na^+^ and 5-HT, preventing reversal of the inward transport step in the reaction cycle (see Fig. 2B).

**Figure 1.**
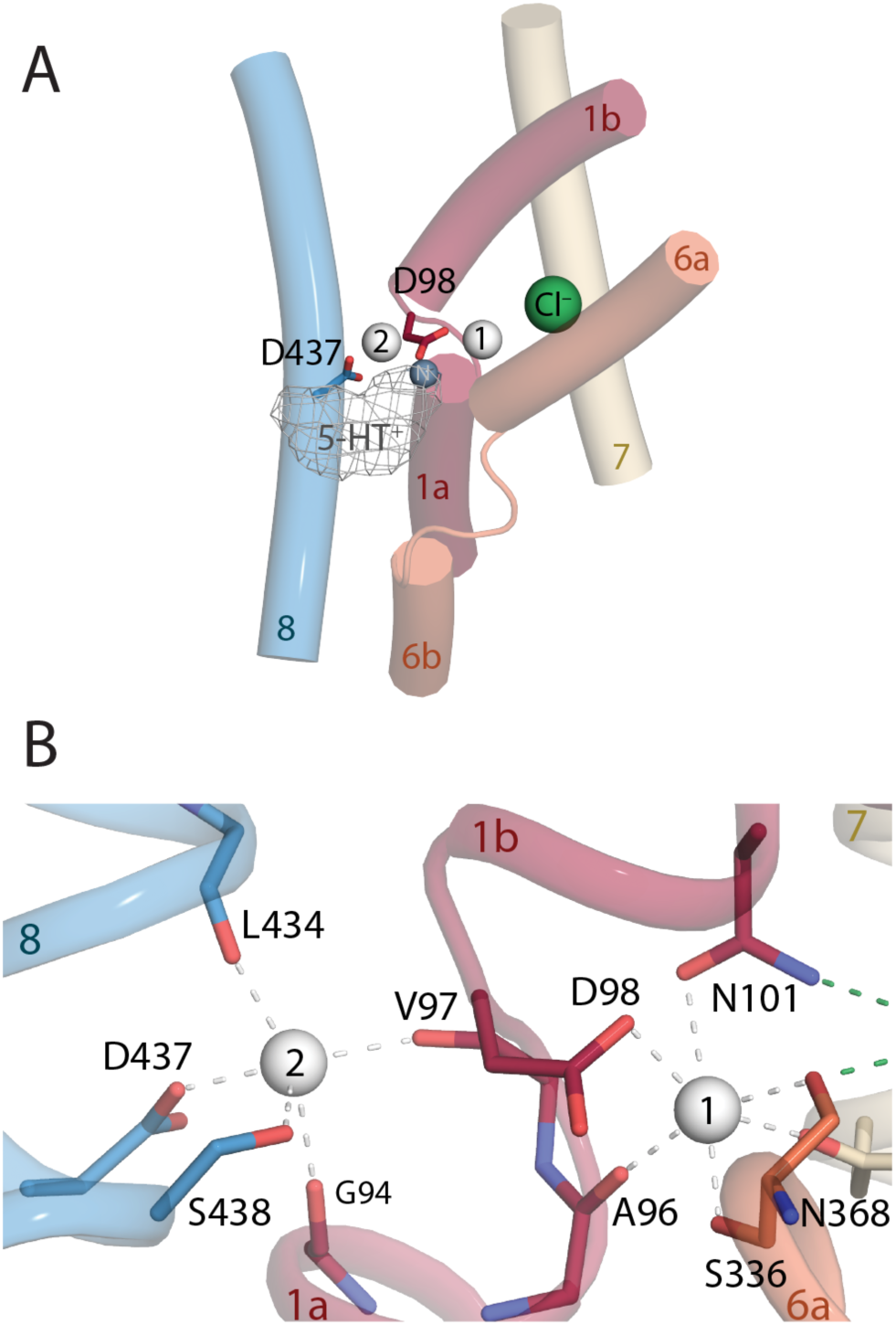
Structure of SERT indicating the possible locations of the K^+^ binding sites tested in this work. (**A**) Overview of the binding site region showing the Ct1 and Ct2 sites (white spheres) and the neighboring chloride ion (green sphere). These binding sites are located close to the interface between the so-called bundle (shades of red/orange) and scaffold (blue/gray) helices. For reference, the so-called S1 substrate (gray mesh) binding site and the primary amine of serotonin (gray sphere) are shown, although the 5-HT molecule itself is not present in the simulations, reflecting the K^+^- driven step that follows 5-HT uptake in the physiological cycle. The protonatable side chains Asp98 in TM1 and Asp437 in TM8 are shown in red and blue sticks, respectively. (**B**) Closeup of the Ct1 and Ct2 sites in the outward-occluded conformation of SERT, with two Na^+^ ions bound, and key coordinating groups shown as sticks.

**Figure 2.**
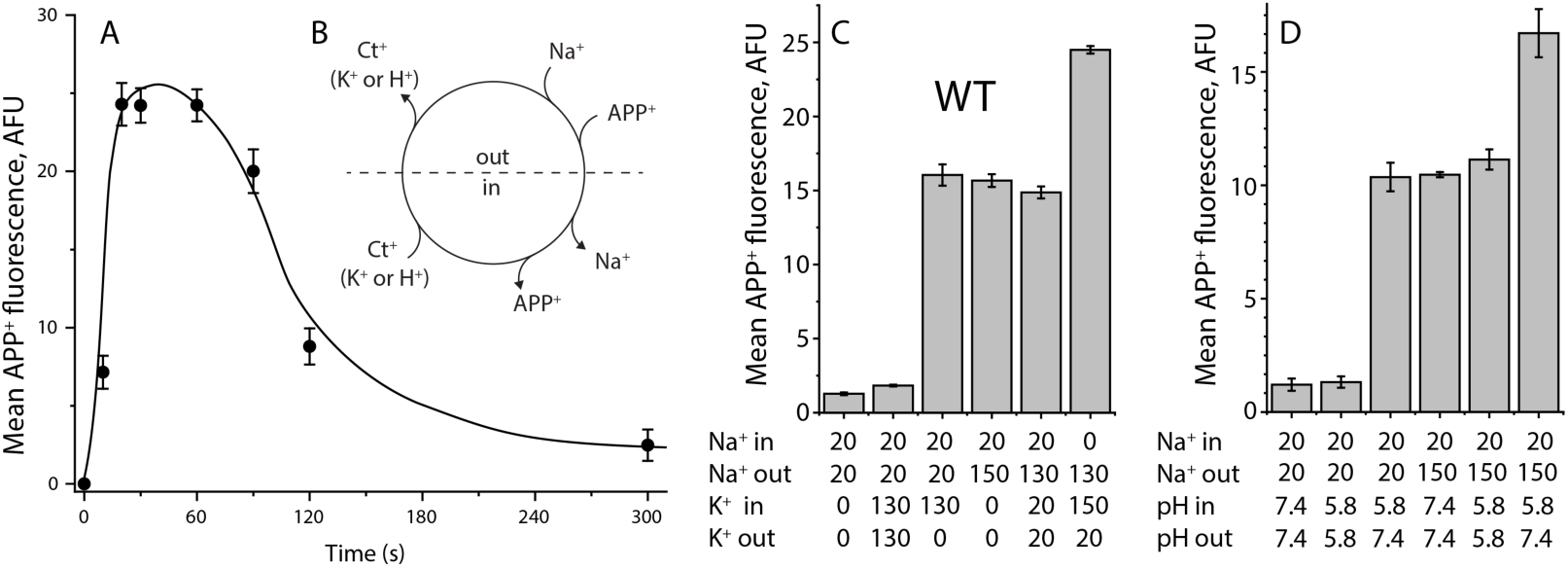
APP^+^ uptake by reconstituted SERT proteoliposomes and influence of intravesicular K^+^ or H^+^ thereon. (**A**) Time course of APP^+^ uptake by reconstituted SERT proteoliposomes. The APP^+^ uptake assay was performed by incubating reconstituted SERT proteoliposomes with 2 μM APP^+^ in an external buffer containing 20 mM NaPi, pH 7.4 and 130 mM NaCl for time intervals of 0–5 min. The proteoliposomes were preloaded with an internal buffer containing 20 mM KPi, pH 7.4 and 130 mM KCl. At each time point, the accumulated APP^+^ in SERT proteoliposomes was measured as described under “Materials and Methods” (n = 3). (**B**) A schematic representation of the APP^+^ transport cycle by SERT proteoliposomes. Ct^+^ indicates a cation that could be either K^+^ or H^+^. (**C**) The influence of intravesicular K^+^ on APP^+^ uptake by SERT WT proteoliposomes. The indicated internal and external buffers were used for reconstitution of SERT proteoliposomes and to assay transport, while maintaining an equal pH (7.4) on both sides of the membrane. NaCl and KCl were supplemented by NMDGCl to maintain the same ionic molarity while generating a Na^+^ inward-directed gradient or/and a K^+^ outward-directed gradient, as indicated. (**D**) The influence of intravesicular proton on APP^+^ uptake. Internal and external buffers containing NaPi and NaCl supplemented by NMDGCl for generating a H^+^ outward-directed gradient or/and a Na^+^ inward-directed gradient, respectively, were used for SERT reconstitution and to assay transport. In these transport assays (C & D), APP^+^ uptake, expressed as accumulated APP^+^ fluorescence in SERT proteoliposomes (mean AFU), was measured by incubating with 2 μM APP^+^ for 1 min at 22°C (n = 3).

SERT adopts the so-called LeuT fold (7), comprising twelve membrane-spanning helices that can be roughly divided into two regions known as the bundle and scaffold. Orientational changes of bundle elements – mainly transmembrane (TM) helices TM1 and TM6 – relative to the scaffold, mediate alternating accessibility of the central binding site (8, 19, 20). Binding of a Na^+^ ion to the so-called Na2 site at the bundle-scaffold interface is critical for mediating the adoption of an outward-facing conformation (21). The Na2 site is proximal to the S1 site (22–24), another Na^+^ site, Na1 (7), and a Cl^−^ binding site (7, 25) (Fig. 1). However, the site of K^+^ binding remains elusive.

Here, we present evidence for the location of the K^+^ ion binding site in SERT using simulations and site-directed mutation studies. The earlier studies of SERT in PPMVs relied on native SERT protein, precluding mutational studies. To interrogate the contributions of individual residues of SERT, we used WT and mutant SERT extracted from a heterologous expression system and reconstituted into liposomes (26). We verify that substrate accumulation by wild type SERT into proteoliposomes can be driven by symport of Na^+^ and antiport of K^+^ or H^+^. Then, to predict the most likely binding site for K^+^, we carried out molecular dynamics (MD) simulations systematically varying the bound cations, as well as the protonation states of two key Asp residues. These simulations reveal the intricate interplay between Na^+^, K^+^ and H^+^ binding in SERT, while predicting that Na2 (hereafter referred to as the second cation site, Ct2) is the most likely site for binding of K^+^ or H^+^. Finally, to test this hypothesis, we measured uptake by proteoliposomes containing SERT modified at the Ct2 site, and the response of those mutants to intracellular K^+^ under whole cell patch clamp. Our data show that K^+^ (or H^+^) accelerates transport by replacing the Na^+^ ion at Ct2 during the return step of the transport cycle and that modification of this site can selectively target the ability of K^+^ or H^+^ to replace Na^+^ in return step of the transport cycle.

## Results

### A K^+^ gradient drives uptake in wild type SERT proteoliposomes

To provide control of the solutions on both sides of the membrane while probing the K^+^ and H^+^ dependence of transport by wild type and mutagenized SERT, we used SERT proteoliposomes reconstituted from membrane patches excised using diisobutylene maleic acid (DIBMA) based on a procedure published previously (26). Uptake of the substrate analog APP^+^ in the presence of 130 mM NaCl was then measured as fluorescence accumulation over time. The APP^+^ accumulation rapidly increased, and then decayed as the ionic driving forces diminished (Fig. 2A). Uptake of substrate by SERT was shown in PPMV to be driven by an inward-directed Na^+^ gradient (see Fig. 2B). Similarly, no uptake into proteoliposomes was observed in the absence of an ion gradient, even in the presence of Na^+^ and K^+^ (Fig. 2C, bars 1 and 2), i.e., with 20 mM Na^+^ on both sides of the membrane. However, APP^+^ accumulation (see Methods) increased significantly upon application of a transmembrane Na^+^ gradient with 150 mM Na^+^ in the external solution (Fig. 2C, bar 4), consistent with observations made with PPMV (4) and this stimulation did not require K^+^ (Fig. 2C, bars 4 vs. 5). To test whether uptake could be driven by high concentrations of K^+^, we applied 130 mM K^+^ to the solution on both sides of the membrane; the level of fluorescence did not increase above background (Fig. 2C, bar 2). By contrast, an outward-directed gradient of K^+^ (in the absence of a Na^+^ gradient) with 130 mM K^+^_in_ and 0 mM K^+^_out_ stimulated uptake to a similar level as the inward-directed gradient of Na^+^ (Fig. 2C, bar 3). Moreover, simultaneous application of an inward-directed Na^+^ gradient and an outward-directed K^+^ gradient elicited an additive effect (Fig. 2C, bar 6). These high levels of uptake depended on the gradient, and not merely on the presence of K^+^ on both sides of the membrane (Fig. 2C, bar 5). Taken together, these results are in strong support of the role of Na^+^:K^+^ exchange during 5-HT^+^ transport as previously measured in PPMV (3, 4).

### An H^+^ gradient drives uptake in wild type SERT proteoliposomes

Although a transmembrane gradient of K^+^ stimulated APP^+^ accumulation, the presence of K^+^ was not required (Fig. 2C, bars 4 vs. 5). Previous studies suggested that H^+^ could compete with K^+^ ions during the antiport phase of the 5-HT uptake cycle and could fulfill the requirement for an antiported cation (11, 18) (Fig. 2B). We therefore tested whether a similar effect was observable in the APP^+^ uptake assay. To establish a baseline, we measured the APP^+^ fluorescence of the proteoliposomes in the presence of 20 mM Na^+^ with the pH on both sides fixed at either 7.4 or 5.8. Neither condition elicited substantial accumulation of APP^+^ (Fig. 2D, bars 1 and 2), consistent with the absence of an ionic driving force. We therefore repeated these measurements in the presence of an outward-directed H^+^ gradient (pH_in_ = 5.8, pH_out_ = 7.4), which was sufficient to drive uptake (Fig. 2D, bar 3) to a similar level as an inward-directed Na^+^ gradient at either pH 7.4 or 5.8 (Fig. 2D, bars 4 and 5, respectively). Similar to the results observed for K^+^ above, the driving force garnered by the outward-directed H^+^ gradient was additive when applied in the presence of an inward-directed Na^+^ gradient (Fig. 2D, bar 6). Overall, these effects echo those obtained for K^+^, supporting previous observations that under appropriate conditions, either K^+^ or H^+^ antiport will drive substrate uptake by SERT.

### MD simulations reveal the preferred K^+^ ion coordination in the known SERT cation sites

While K^+^ antiport by SERT is firmly established, the location of the binding site for K^+^ is unknown. Since H^+^ can substitute for K^+^ ions, the binding site is likely to contain one or more protonatable side chains. Consequently, two plausible locations for K^+^ binding include the Na1 and Na2 sites, each of which contains one Asp side chain, namely Asp98 and Asp437, respectively (Fig. 1). Ct1 and Ct2 (Fig. 1B) are highly conserved across the wider neurotransmitter:sodium symporter family (27), having first been reported in the eubacterial transporter LeuT (19). The Ct1 site is adjacent to the Cl^−^ ion site (Fig. 1A) and this proximity is likely responsible for the inter-dependence of their binding affinities (25, 28). However, despite the conservation of both Ct1 and Ct2 in SERT, the stoichiometry of 5-HT^+^ to Na^+^ is 1:1 (3), raising the possibility that, like the Cl^−^ ion, one of the two Na^+^ ions may be bound also during the “return” step of the cycle (13). To assess whether either of the two known Na^+^ sites is suitable for K^+^ binding, we carried out MD simulations of the substrate-free transporter (Fig. S1). We assumed simultaneous occupancy of one Cl^−^ ion, one Na^+^ ion, and one K^+^ ion (in either of the two sites), while also assessing all combinations of protonation states of the two Asp side chains.

Because K^+^ enters from the cytoplasm during the physiological transport cycle, we first attempted to assess the binding of Na^+^ and K^+^ to SERT structures in which the cytoplasmic pathway is open. However, this approach proved problematic due to the frequency of unbinding and binding events (measured as distance to the ion binding residues; Fig. S2A), reflecting in part that the (K^+^) sites are not suitably organized, due to the separation of TM1 and TM8 in that state; moreover, the pathway is not expected to close reproducibly on a micro-second timescale even in the presence of K^+^. By contrast, simulations of an occluded conformation of substrate-free SERT produced trajectories in which the ion behavior was reproducible over repeated simulations on a 500 ns timescale (Fig. 3, S3).

**Figure 3.**
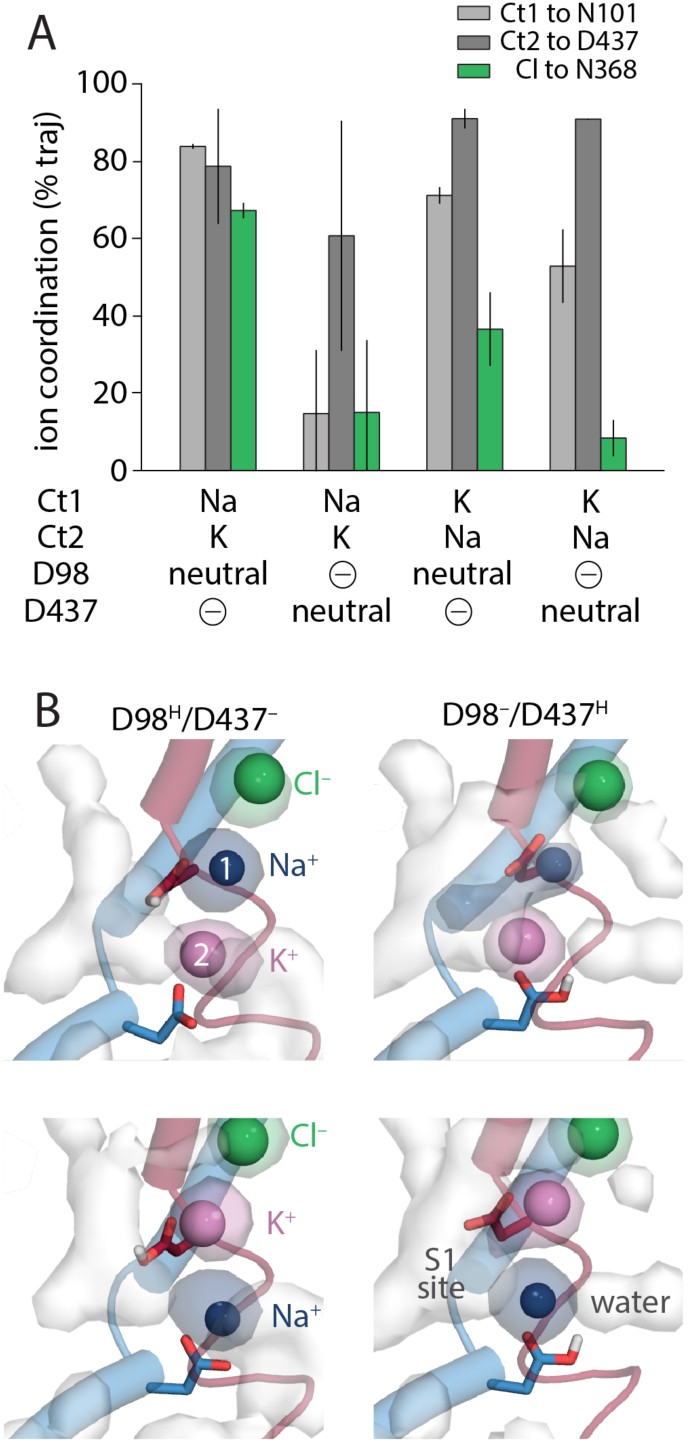
Stability of cation and Cl^−^ coordination as a function of the bound alkali ions and protonation state of the two key binding site aspartate side chains in outward-occluded conformation of SERT in the absence of 5-HT. (**A**) Coordination is defined using the distance to a representative side chain (N101, D437 or N368) in each of the three ion binding sites (Ct1, Ct2, and chloride, respectively), and plotted as percentage of trajectory length, with error bars indicating the standard deviation over n=4 simulations. An ion was considered to be coordinated if its distance to the representative side chain non-hydrogen atom(s) was below a threshold of 2.6 Å, 2.9 Å or 3.6 Å for Na^+^, K^+^ and Cl^−^ ions, respectively. The configuration in which D98 is protonated and D437 is charged, with Na^+^ in Ct1 and K^+^ in Ct2 (leftmost bars) is predicted to have the most stable ion coordination overall. Corresponding time courses are shown in Fig. S3. (**B**) Ion and water occupancy during MD simulations. Densities are shown for simulations in which cation site 1 (Ct1) is occupied with either Na^+^ (upper panel) or K^+^ (lower panel), while cation site 2 (Ct2) is occupied with either K^+^ (upper panel) or Na^+^ (lower panel). Columns differ by the position and number of protons that are covalently bound to the binding site asp residues, as indicated. Occupancy maps (surfaces) are averages over repeated simulation trajectories, with water in white, Na^+^ in dark blue, K^+^ in violet, and Cl^−^ in green. Similar data is shown for all simulation systems in Fig. S4.

We assumed that stable coordination, measured as the distance between the ion and its (representative) binding site residues in the occluded conformation, reflects a thermodynamically (meta)stable intermediate, consistent with previous observations of Na^+^ binding in occluded conformations of LeuT-fold transporters on a sub-microsecond timescale e.g. (29, 30). Analysis of the ion-protein distances revealed a striking dependence on the protonation state of the coordinating Asp groups. In particular, when both Asp were charged, the ions were poorly coordinated (Fig. S3) and poorly localized to the sites (Fig. S4). By contrast, consistency in the coordination and position was observed in simulations in which one of the two aspartates was protonated. Of these, the combination with the smallest perturbations in coordination (Fig. 3A) and position (Fig. 3B) consisted of a Na^+^ ion at Ct1, coordinated in part by Asp98 in its protonated form, and a K^+^ ion at Ct2, coordinated in part by Asp437 in its charged form.

Significant perturbations were observed in all simulations with a K^+^ ion placed at Ct1. In particular, the Cl^−^ ion coordination was influenced strongly by the character of the neighboring cation (Fig. 3; Fig. S4; S5), which affects the anion interactions with Asn368 (Fig. S6) and allows water penetration into the Cl^−^ site (Fig. 3B; Fig. S4). Thus, the simulations predict that K^+^ binds at Ct2, while a constitutive Na^+^ ion preferentially occupies Ct1. Notably, in this scenario, Asp98 is predicted to be protonated because that is the most stable configuration, unlike in the 5-HT bound state where the charged primary amine on the substrate, we believe, is directly interacting with Asp98 in its charged state.

### MD simulations with a K^+^ ion in S1 support the preferred ion combination

In the aforementioned simulations, the S1 site is occupied by water (Fig. 3B). Although to some this might be unexpected, such hydration is not an inherently unstable occurrence, as it is evidently preferable to a vacuum. Nevertheless, the presence of water in this region raises the possibility that K^+^ ions compete with 5-HT by occupying the same location as the substrate primary amine group, which shares Asp98 coordination with Na1 (Fig. 1A). If the K^+^ ion is in the S1 site, and we assume that only one Na^+^ and one Cl^−^ ion are bound, as above, then either Ct1 or Ct2 must be unoccupied. Given its limited impact on the Cl^−^ ion, we assumed that the constitutive Na^+^ ion would occupy the Ct1 site. We carried out MD simulations of SERT with a K^+^ ion initially placed at the amino nitrogen of 5-HT^+^ next to Asp98.

Tracking the position of the K^+^ ion relative to the two sites demonstrates again an exquisite sensitivity to the protonation states of the aspartate residues (Fig. 4). In particular, when Asp98 was protonated and Asp437 was charged (Fig. 4, leftmost column), the K^+^ ion placed in the S1 site rapidly and reproducibly moved away from the neutralized Asp98 (Fig. 4, upper panel) and into the Ct2 site toward the charged Asp437 side chain (Fig. 4, lower panel; Movie S1). The endpoint of these trajectories was the configuration identified in the previous section, i.e., with Na^+^ at Ct1, K^+^ at Ct2, and Asp98 protonated. By contrast, when Asp98 was charged, or when both Asp were protonated, the K^+^ ion remained in the central pocket (Fig. 4, Movie S2), as if encountering a kinetic barrier to entering the Ct2 site, presumably due to electrostatic interaction between the K^+^ ion and the charge on Asp98. Notably, the observed movement into Ct2 could not be attributable purely to attraction to the charge at Asp437, because no similar transitions occurred when both Asp were charged (Fig. 4, rightmost column). When in the S1 site the K^+^ ion formed interactions that alternated between the backbone of Phe335 and the side chain of Ser438, in addition to a cation-pi interaction with the aromatic ring of Tyr95 (Fig. S7). However, this coordination was overall less stable than that observed when K^+^ occupied Ct2 and Asp98 was protonated (Fig. 4). Thus, taken together with the results above, the simulations also predict that K^+^ binds at Ct2.

**Figure 4.**
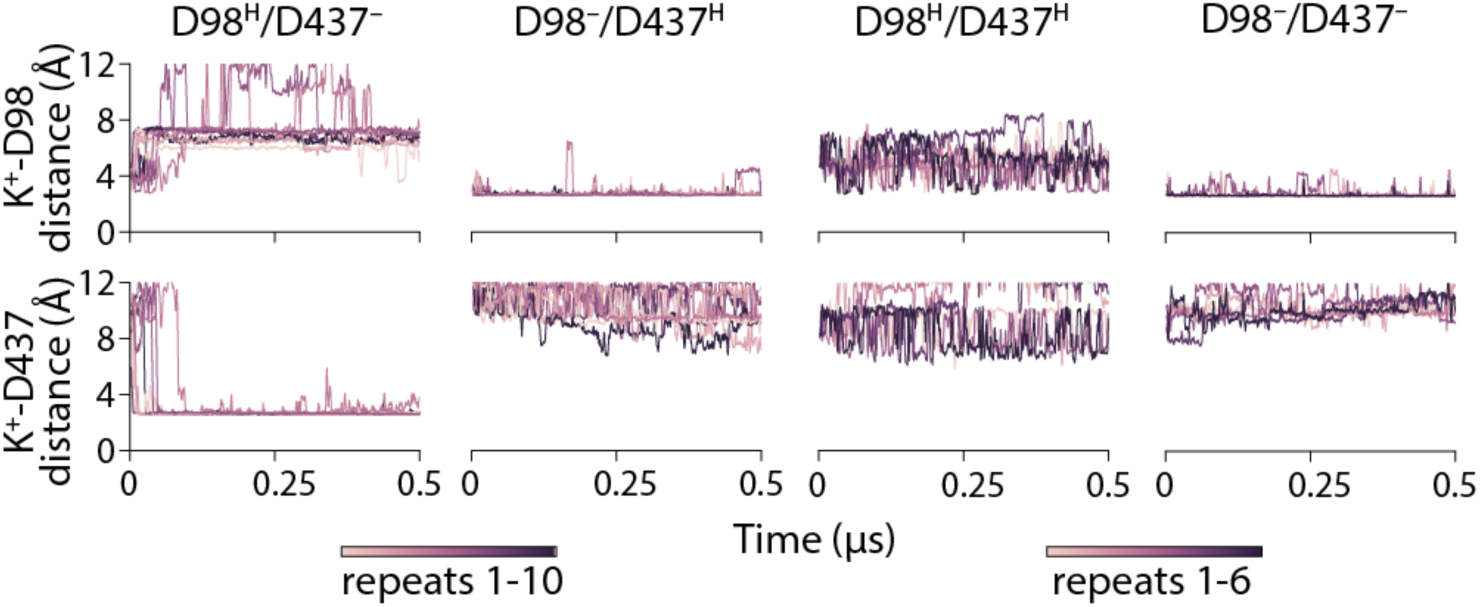
K^+^ enters cation site 2 starting from the 5-HT^+^ amino position only under specific protonation states. Ion distances were measured to the side-chain oxygen atoms of either Asp98 (upper panel) or Asp437 (lower panel), using the minimum of the two oxygen atoms at each frame. Columns indicate the simulations carried out with different protonation states: Asp98 protonated (far-left- and center-right columns) or Asp437 protonated (center columns). Lines indicate individual trajectories: n = 10 for singly-protonated systems and n = 6 for fully charged or doubly-protonated systems, see scale bars.

### The substrate-induced current through D437T and S438A is insensitive to K^+^_in_

To test the hypothesis that K^+^ binds at Ct2, we made SERT mutants where we either replaced Asp437 with Thr (to recapitulate the binding site present in LeuT) or replaced Ser438 with Ala (see Fig. 1B). We then recorded 5-HT^+^-induced currents obtained from cells transiently expressing these mutants. For comparison, we first replicated measurements made for wild type SERT (11, 15) at a range of applied voltages (Fig. 5A, 5B). Upon application of 5-HT under physiological intracellular K^+^ concentrations, wild type SERT rapidly elicited an inward peak current (Fig. 5A), reflecting the movement of 5-HT and Na^+^ charges across the membrane for all the transporters that were poised in an outward-facing conformation, and the subsequent release of Na^+^ ions into the cytosol (Fig. 5K) (11). The peak current was also observed when the intracellular solution contained NMDG^+^ instead of K^+^ (Fig. 5B). As shown previously (11), the magnitude of the initial inward current is linearly dependent on voltage; however, the slope of this dependence becomes shallower if K^+^ is present in the intracellular solution (Fig. 5C), because intracellular K^+^ (or Na^+^) offsets the charge movement by binding to the inward-facing conformation (Fig. 5K).

**Figure 5.**
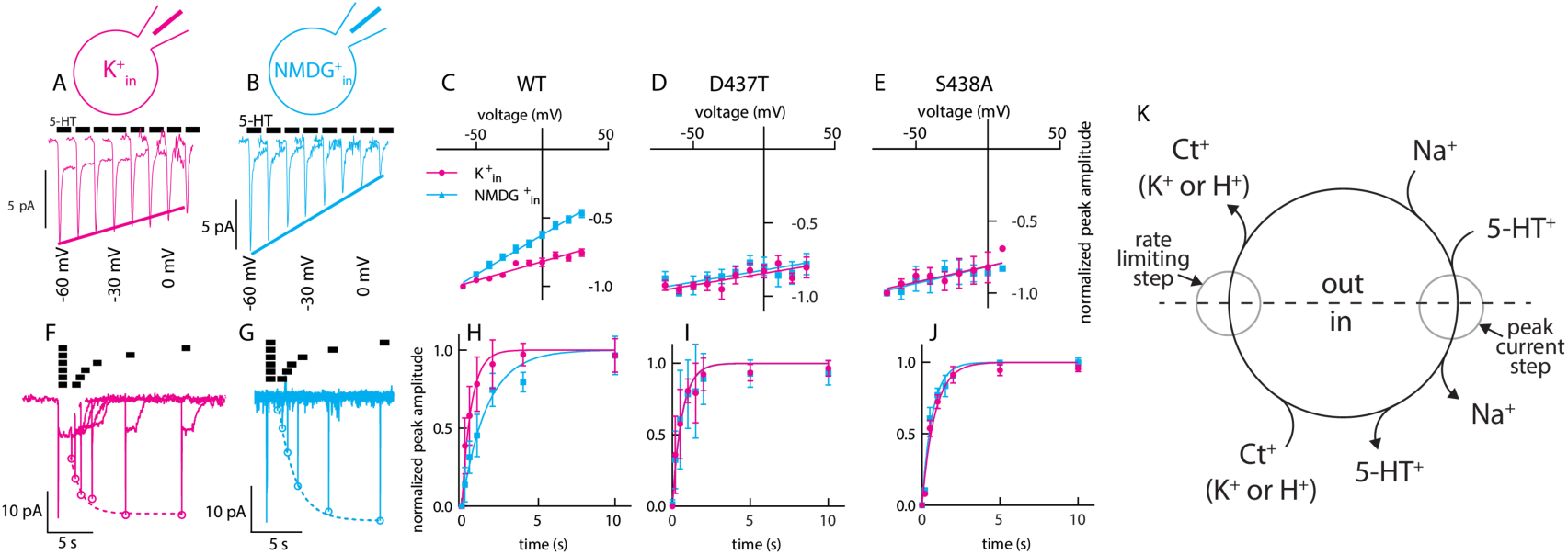
The substrate-induced current through D437T and S438A is insensitive to K^+^_in_. (**A, B**) Representative traces of peak currents obtained from wild type SERT in the presence of 160 mM K^+^_in_ (A, magenta) or 160 mM NMDG^+^_in_ (B, cyan) recorded at voltages ranging from –60 mV to +10 mV. (**C**) Plot of the normalized peak current amplitudes measured from wild type SERT as a function of voltage in the presence and absence of 160 mM K^+^_in_ (K^+^ n = 7, magenta circles, NMDG^+^ n = 15, cyan squares). The magenta and cyan lines are linear fits to the data points. The slope was 2.78 ± 0.22 V^-1^ and 5.81 ± 0.26 V^-1^ for 160 mM K^+^_in_ and NMDG^+^_in_, respectively. (**D**) The same as in C for SERT D437T (K^+^ n = 14, circles, NMDG^+^ n = 15, squares). The slope was 1.59 ± 0.49 V^-1^ and 1.63 ± 0.56 V^-1^ for 160 mM K^+^_in_ and NMDG^+^_in_, respectively. The corresponding traces are shown in Fig. S8A and S8B. (**E**) The same as in C for SERT S438A (K^+^ n = 7, circles, NMDG^+^ n = 8, squares). The slope was 2.22 ± 0.9 V^-1^ and 2.36 ± 0.9 V^-1^ for 160 mM K^+^_in_ and NMDG^+^_in_, respectively. The corresponding traces are shown in Fig. S8C and S8D. (**F**) Schematic representation of the transport cycle of SERT. Indicated in this scheme are the binding and unbinding reactions of the (co)-substrates, the transition that gives rise to the peak current, and the rate-limiting step. (**G, H**) Representative current traces resulting from a two-pulse protocol recorded from wild type SERT. The intracellular solution contained 160 mM K^+^ (G) or 160 mM NMDG^+^ (H). (**I**) Plot of the normalized peak current amplitudes of wild type SERT as a function of the time interval between the first and the second pulse (K^+^ n = 11, circles, NMDG^+^ n = 10, squares). The peak current recovery rate, as estimated by a monoexponential fit to the data points, was 1.64 ± 0.12 s^-1^ and 0.61 ± 0.04 s^-1^ in the presence of 160 mM K^+^_in_ and NMDG^+^_in_, respectively. (**J**) The same as in I but for SERT D437T (K^+^ n = 5, circles, NMDG^+^ n = 5, squares). The peak current recovery rate was 1.61 ± 0.19 s^-1^ with 160 mM K^+^_in_ and 1.57 0.22 s^-1^ with 160 mM NMDG^+^_in_. The corresponding traces are shown in Fig. S8E and S8F. (**K**) The same as in J but for SERT S438A (K^+^ n = 7, circles, NMDG^+^ n = 6, squares). The peak current recovery rate was 1.21 ± 0.1 s^-1^ with 160 mM K^+^_in_ and 1.51 ± 0.12 s^-1^ with 160 mM NMDG^+^_in_. The corresponding traces are shown in Fig. S8G and S8H.

Having recapitulated earlier observations for wild type SERT, we then asked whether the amplitude of the peak current through D437T and S438A was modulated by K^+^_in_. Comparing current amplitudes with and without K^+^_in_ revealed that the voltage dependence of the peak current amplitude was unaffected by K^+^_in_ in both D437T (Fig. 5D; Fig. S8A-B) and S438A (Fig. 5E; Fig. S8C-D). Thus, the transport-associated current carried by the Ct2-site mutants was no longer sensitive to the presence of K^+^_in_.

### Substrate-dependent turnover rates by D437T and S438A are insensitive to K^+^_in_

In wild type SERT, the substrate turnover rate was shown to be accelerated in the presence of K^+^_in_, by using a two-pulse electrophysiological measurement. In this experiment, the first application of 5-HT^+^ (i.e., 30 µM) enables all transporters to enter the transport cycle. The time course of recovery to the resting state (i.e., the Na^+^-bound, outward-facing state) is then monitored by applying a second pulse of 5-HT^+^ at different time intervals after the first pulse. Because the transporters can return to the resting state only after completing a full cycle, the rate of recovery of the peak current reflects the substrate turnover rate, whose rate-limiting step is controlled by the intracellular cation binding (Fig. 5K). Comparing data obtained for wild type SERT in the presence of 160 mM K^+^_in_ (Fig. 5F) with that obtained in the absence of K^+^_in_ (Fig. 5G), plotted as a function of the time interval (Fig. 5H), revealed that the transporter turnover rate increased from 0.62 ± 0.06 s^-1^ in the presence of 160 mM NMDG_in_^+^ to 1.79 ± 0.19 s^-1^ in the presence of 160 mM K^+^_in_, in agreement with Bhat et al. (15).

To test whether the K^+^-dependence of the recovery rate involves the Ct2 site, we repeated these recordings for cells expressing D437T (Fig. 5I and Fig. S8E-F) and S438A (Fig. 5J and Fig. S8G-H). In both cases, the dependence on the pulse interval was essentially the same in the absence and presence of K^+^_in_. The failure of K^+^_in_ to accelerate substrate uptake is consistent with the idea that K^+^ cannot bind to SERT when the Ct2 site is compromised.

### The leak current through Ct2 site mutants is insensitive to K^+^_in_

In the presence of internal K^+^, wild type SERT exhibits a steady inward current for the duration of 5-HT application (Fig. 6A), which is not observed in when intracellular K^+^ is replaced by NMDG^+^ (Fig. 6D) (14, 31). This steady inward current involves a conductive intermediate formed during the return step in the reaction cycle (Fig. 6J). If formation of the conductive intermediate requires binding of K^+^ at the Ct2 site, this steady current should be abolished in the Ct2-site mutants. Indeed, currents measured at –10 mV for both D437T (Fig. 6B) and S438A (Fig. 6C), lack the steady inward current observed for WT at the same voltage (Fig. 6A). The data in Fig. 6 is restricted to currents recorded at –10 mV because of an additional leak current in these mutants that interferes with the measurements at more negative potentials (Fig. S9).

**Figure 6.**
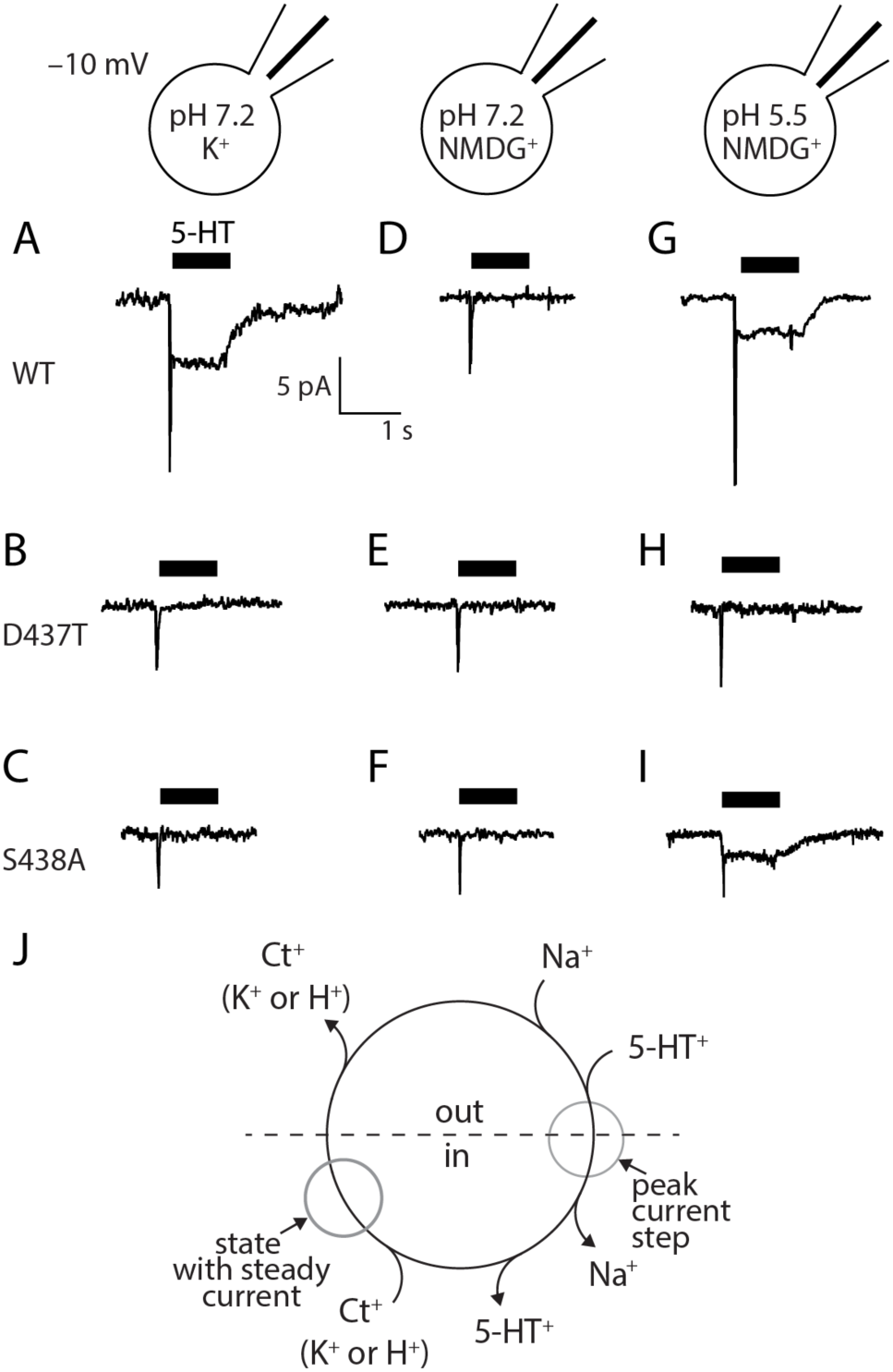
Dependency of SERT steady currents on K^+^ and H^+^ in Ct2-site mutants. (**A-C**) Representative current induced by 30 µM 5-HT recorded from wild type SERT (A), D437T (B), and S438A (C) at –10 mV in the presence of 160 mM K^+^_in_ and with the intracellular pH set to 7.2. (**D-F**) Representative current induced by 30 µM 5-HT recorded from wild type SERT (D), D437T (E), and S438A (F) at –10 mV in the presence of NMDG mM K^+^_in_ and with the intracellular pH set to 7.2. (**G-I**) Representative current induced by 30 µM 5-HT recorded from wild type SERT (G), D437T (H), and S438A (I) at –10 mV in the presence of NMDG mM K^+^_in_ and with the intracellular pH set to 5.5. (**J**) Schematic of the transport cycle of SERT. Highlighted in this scheme are the steps in the transport cycle where the peak current and the steady current are produced.

### Accumulation by Ct2-site mutants is unaffected by a K^+^ gradient

Given that Ct2-site SERT mutants lacked sensitivity to intracellular K^+^ in their conductance properties, we asked whether APP^+^ accumulation by these mutants would also differ. Similar to wild type SERT (Fig. 2C), no substrate accumulation was observed in the absence of a Na^+^ gradient, both with and without high concentrations of K^+^ on both sides of the membrane for D437T (Fig. 7A, bars 1 and 2). Nevertheless, D437T was still capable of uptake: the presence of a Na^+^ gradient drove APP^+^ uptake that was unaffected by 20 mM K^+^_in=out_ (Fig. 7A, bars 4 and 5; Table S1). Unlike wild type SERT however, the presence of an outward-directed K^+^ gradient alone did not result in APP^+^ accumulation in D437T (Fig. 7A, bar 3), nor did the addition of an outward-directed K^+^ gradient produce an additive effect on uptake driven by a Na^+^ gradient (Fig. 7A, bar 6). Similar observations were made for S438A (Fig. 7C, Table S1). The dashed bars in this figure indicate the accumulation with WT SERT, normalized to the accumulation with a Na^+^ gradient alone. Thus, D437T and S438A are insensitive to K^+^ gradients, suggesting that the wild type protein binds K^+^ at the Ct2 site, as predicted by the simulations and that this site is essential for a transmembrane K^+^ gradient to contribute to substrate accumulation. We also tested D437A, but it was not active, consistent with the requirement for a polar group to bind Na^+^ during transport (Table S1).

**Figure 7.**
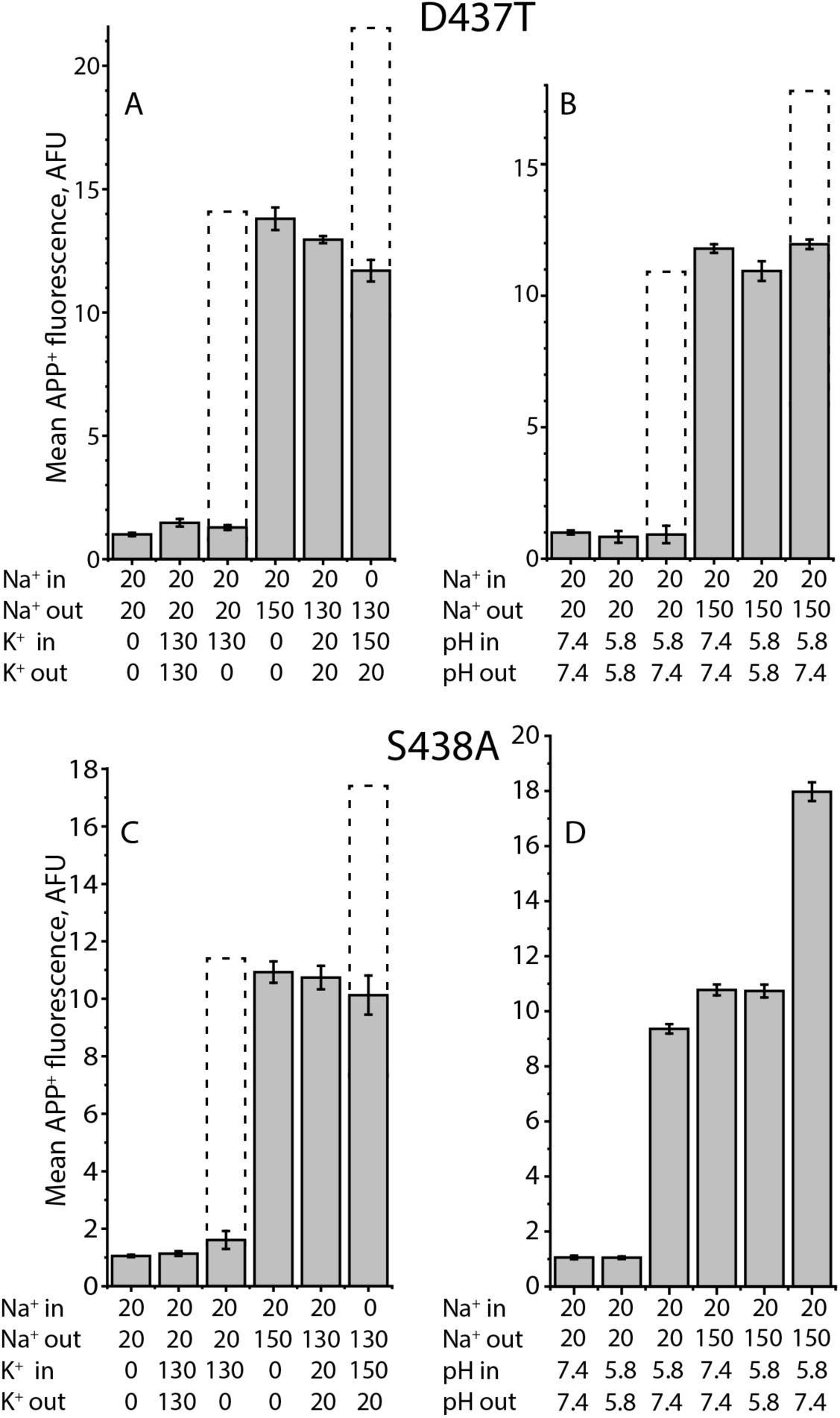
The effect of substitutions of Ct2-site residues on the ability of intravesicular K^+^ or H^+^ to stimulate APP^+^ uptake. (**A, B**) The effect of substitution of Asp437 with Thr on the stimulation of APP^+^ uptake by intravesicular K^+^ (A) or H^+^ (B). (**C, D**) The effect of substitution of Ser438 with Ala on the stimulation of APP^+^ uptake by intravesicular K^+^ (C) or H^+^ (D). In these transport assays, various combinations of internal and external buffers were used for reconstitution of D437T, or S438A proteoliposomes and to assay APP^+^ uptake, as indicated in the legend to Figure 1 (n = 3). Dashed bars indicate the accumulated APP^+^ fluorescence values in the WT SERT proteoliposomes assayed in parallel and in the same conditions.

### H^+^-driven accumulation of substrate requires Asp437

Given that a H^+^ gradient can replace a K^+^ gradient in wild type SERT (18) (Fig. 2B, 2D), we expected that if D437 is also the sought-after H^+^ binding site, a pH gradient would be effective in S438A, but not in D437T. We therefore measured APP^+^-dependent fluorescence in liposomes containing D437T and S438A under various pH conditions. Unlike wild type SERT, D437T was unresponsive to the outward-directed H^+^ gradient (Fig. 7B, bar 3), as well as to the additive effect in the presence of an inward-directed Na^+^ gradient (Fig. 7B, bar 6). Notably, D437T retained Na^+^ dependence and was not affected by pH changes when the pH was the same on both sides (Fig. 7B, bars 4 and 5).

The S438A mutant behaved similarly to wild type; it responded to imposition of an outward-directed H^+^ gradient (Fig. 7D, bar 3) and that response was additive with the presence of an inward-directed Na^+^ gradient (Fig. 7D, bar 6). Taken together, these results suggest that an H^+^ ion can continue to bind at Asp437 when Ser438 is removed, but also that the ionizable group at Asp437 is critical for H^+^-driven uptake by SERT.

### The H^+^-dependence of SERT requires Asp437

In patch clamp recordings of SERT, H^+^ can also functionally replace K^+^_in_ (11). Specifically, in the absence of K^+^_in_, lowering the intracellular pH from 7.2 to 5.5: (i) doubled the rate of substrate uptake; (ii) restored the substrate-induced steady current; and (iii) altered the voltage dependence of the peak currents. To test whether Asp437 is the H^+^ binding site, we recorded substrate-induced currents carried by D437T and S438A in the absence of intracellular K^+^, but in the presence of high and low H^+^_in_ (i.e., pH 5.5 and 7.2, respectively) (Fig. 6D-I). As for proteoliposome uptake, we posited that a low intracellular pH ought to be effective in S438A but not in D437T. This was indeed the case. At pH 5.5, D437T exhibited no steady current during the application of 5-HT (Fig. 6H), unlike both the wild type SERT (Fig. 6G) and S438A (Fig. 6I).

Measurements of 5-HT-elicited peak currents at a range of voltages were consistent with these findings (Fig. S10). For D437T, increasing the intracellular H^+^ concentration had minimal impact on the current-voltage relationship (Fig. S10A-C), indicating that the removal of the acidic side chain modifies the response of SERT to low pH_in_ (11). For S438A, however, the currents were sensitive to the intracellular pH (Fig. S10D-F). Specifically, intracellular H^+^ in S438A can mimic the role of intracellular K^+^, just as in wild type SERT (11). Thus, in S438A, where the aspartate is present, features of the pH dependence of WT SERT are preserved. Taken together, these data show that: (i) the aspartate at position 437 can serve as a binding site for a H^+^; and (ii) the pK_A_ of this residue is voltage dependent.

## Discussion

A dependence on intracellular K^+^ is a eukaryotic adaptation that evolved in members of the SLC6 family (5) as well as in SLC1 transporters (32, 33). This dependence confers an additional cellular strategy for modulating the rate of transport. However, the small difference in radii between the alkali cations raises the question of how K^+^ effectively competes with serotonin intracellularly, while not being co-transported alongside it, e.g., by replacing Na^+^. Understanding the nature of that competition requires identification of its binding location.

In this work, we predicted the most likely site at the molecular level using MD simulations of SERT. Of the two structurally-resolved cation sites, the Ct2 site has been posited to be the K^+^ site (15), consistent the fact that Ct2 (i) lies closest to the cytoplasmic pathway, (ii) is significantly altered during pathway opening (8, 20) facilitating Na^+^ release (34, 35) and (iii), relatedly, is responsible for Na^+^-mediated stabilization of the outward-facing state in related transporters, including LeuT (21). Nevertheless, simulations of the inward-facing state revealed stochastic unbinding of the Na^+^ at the Ct1 site (Fig. S2), raising the possibility that it could be displaced by K^+^. Among the 26 tested cation combinations in the outward-occluded state, those with K^+^ at the Ct1 site showed significant perturbation of both the Ct1 and chloride sites (Fig. S5), and K^+^ ions at the central S1 site were poorly coordinated (Fig. S7). By contrast, K^+^ bound at the Ct2 site was much better accommodated; indeed, binding to Ct2 from S1 occurred spontaneously (Fig. 4).

Replacement of the protonatable Ct2 residue by the side chain present in K^+^-independent bacterial transporters in D437T abolished the behavior reported previously to depend on intracellular K^+^, as did removal of the other Ct2-site side chain at position 438. Notably, these two mutations differed in their responses to intracellular H^+^, providing strong evidence that both residues are required for K^+^ antiport, but only Asp437 is essential for H^+^ antiport. Both Asp437 and Ser438 are conserved across all 87 annotated SLC6A4 sequences in the UniprotKB database as of October 2023. The closely related transporters for dopamine and norepinephrine, DAT and NET, also contain these residues. Indeed, DAT also binds K^+^ ions (15, 16) – presumably at the same site – although it remains to be fully established whether DAT can translate binding into a driving force for dopamine accumulation (17).

If the Ct2 site is their binding site, how then do H^+^ and K^+^ ions compete with Na^+^ and 5-HT? Differences in physiological cytoplasmic concentrations are sufficient to explain how K^+^ competes with Na^+^ for the Ct2 site. However, given that configurations predicted in the simulations with K^+^ at Ct2 are not markedly different from that of the Na2-bound state, it remains to be understood from a molecular standpoint, why 5-HT does not bind together with K^+^ and become exported, i.e., how the transporter prevents futile efflux of substrate. In this context, an important consideration may be the additional stabilization conferred by the protonation of Asp98, directly competing with 5-HT.

When the amino acid transporter LeuT adopts an apo occluded conformation, the S1 site is partially occupied by the side chain of the highly-conserved residue Leu25 (36). We therefore considered whether a similar mechanism is at play in SERT to compensate for 5-HT binding; however, the rearrangement of Leu25 in LeuT requires a major backbone conformational change including of the preceding residue Gly24. In monoamine transporters, by contrast, the equivalent leucine (Leu99 in SERT), is flanked by Asp98, rearrangement of which is more sterically constrained. Thus, competition by protonation of Asp98 is a more likely scenario.

If Asp98 protonation accompanies K^+^ antiport, one might expect the D437T mutant – which preserves Asp98 – to be driven by an outward-directed proton gradient acting at Asp98. Our measurements do not suggest this is the case (Fig. 7B). Although we do not have a definitive explanation for this inconsistency, one possibility is that the requirement for the proton on Asp98 is inextricably linked to the presence of the two protonatable sites, because of direct electrostatic interactions between Asp437 and Asp98, whose side chains can be <7Å apart. It will be of interest in future work to explore the role of protons in accompanying the return step of transport, and thereby in defining the net stoichiometry of transport, in more detail. The identification of Ct2 as the binding site for K^+^ contributes to the groundwork required for such studies in SERT.

## Materials and Methods

### Reconstitution of SERT proteoliposomes

Detergent-free reconstitution of SERT proteoliposomes was performed as described previously (26), but with the following modification. In brief, confluent HEK293T cells stably expressing hSERT wild type (UniprotKB identifier: P31645) or mutant with a 10-His epitope at its C-terminus were collected and disrupted by 3 × sonication using an ultrasonic homogenizer (Scientz, Ningbao, China) in ice-cold lysis buffer containing 50 mM Tris-HCl, pH7.4, 150 mM NaCl, 0.1 mM PMSF, 10% glycerol, 1 mM EDTA, and 0.1% proteinase inhibitor mixture. After removal of cell debris by centrifugation at 10,000 × g for 5 min, cell membrane fraction was collected by re-centrifugation at 100,000 × g for 1 h at 4°C and solubilized in DIBMA solubilizing buffer containing 50 mM Tris-HCl, pH 7.4, 150 mM NaCl, 0.1 mM PMSF, 2.5% DIBMA, and 10% glycerol with a gentle rotation overnight at 4°C. SERT was then purified by a successive 4-step purification procedure, which included incubation with Ni-NTA agarose for 3 h at 4°C, 3 × wash with washing buffer (50 mM Tris-HCl, pH 7.4, 150 mM NaCl, 10% glycerol, and 5 mM imidazole), elution with elution buffer (50 mM Tris-HCl, pH 7.4, 150 mM NaCl, 10% glycerol, and 300 mM imidazole), and Sephadex G-200 size exclusion chromatography to remove imidazole.

The lipid mixture used for reconstitution contained a ratio of 20:40:25:15 (w/w) for CHS (cholesterol), DOPC (1,2-dioleoyl-sn-glycero-3-phosphocholine), DOPE (1,2-dioleoyl-sn-glycero-3-phosphatidylethanolamine), and DOPG (1,2-dioleoyl-sn-glycero-3-[phospho-rac-(1-glycerol)]) (Avanti Polar Lipids) respectively. This mixture was suspended in 50 mM Tris-HCl buffer, pH 7.4, sonicated until translucent at a concentration of 15 mg/mL, diluted into 20 × volumes of an internal buffer, as indicated, and subjected to 3 × repeated freeze-thaw steps. After application of 30 × extrusion using an Avanti mini extruder, the unilamellar vesicles equilibrated with an internal buffer were collected. The purified SERT was then added for a protein/lipid ratio of 1:100 (w/w), vortexed for 1 min, and incubated for 20 min on ice.

### Transport assay with SERT proteoliposomes

To measure transport by reconstituted SERT proteoliposomes, APP^+^ uptake was initiated by diluting 20 μL of SERT proteoliposomes preloaded with an internal buffer into 380 μL of an external buffer containing 2 μM APP^+^. After incubation for time intervals of 0–5 min at 22°C, the reaction was quenched by a rapid addition of 500 μL of ice-cold external buffer. APP^+^ remaining in solution was removed by filtration and the accumulated APP^+^ was measured by the fluorescence of the filtrate. All liposome-based APP^+^ uptake experiments were performed at least three times. Nonspecific uptake was measured in the presence of 10 μM fluoxetine.

### Molecular Dynamics Simulations

Simulations of two conformations of SERT were set up according to a previously-developed protocol (37). The inward-open conformation structure (PDB identifier 6DZZ (8) was truncated at residues 78-85 with Coot v 0.8.9.3-pre (38) due to inclusive densities in the EM map (EMD-8943)), while the outward-occluded conformation was based on a previously-published hybrid model (23). In each case, the protein structure was placed in a hydrated palmitoylolyeoylphosphatidylcholine (POPC) lipid bilayer at a salt concentration of 0.15M NaCl, which was then equilibrated at coarse-grained resolution using Gromacs v2018.8 (39) and the Martini v2.2 force field (40) for 50 µs. To determine the appropriate box size, we analyzed the perturbations of the membrane, which were minimally impacted by the periodic boundary images at a size of 12x12x11 nm. A representative snapshot with minimal structural deviation from the equilibrated bilayer was converted to an all-atomistic system by backmapping (41) of lipid, water, and ions. The protein was replaced by the all-atomistic model and cavities in the transporter were solvated with Dowser (42). The fully-atomistic simulations were run with NAMD v2.13 or v3.0a9 using the CHARMM36m force field (43) with TIP3P waters (44). Residue Glu508 was set to be protonated, based on proximity to Glu136 (45), and residues C200 and C209 were disulfide bridged. The protonation states of residues Asp98 and Asp437 were varied to cover all possible combinations. In the outward-occluded conformation, Ct1 and Ct2 contained either two Na^+^ ions, two K^+^ ions, or mixtures of the two cation types. The only exception was a condition with a Na^+^ ion in Ct1, Ct2 empty, and a K^+^ ion placed in the S1 binding site at the position of the primary amine N of ibogaine (8). In the inward-open conformation, Ct1 and Ct2 contained either two Na^+^ ions, one Na^+^ at Na1, or no cations. The periodic box contained ∼150,000 atoms with dimensions of ∼11 nm^3^ after equilibration.

Protein and dowser water hydrogens were energy-minimized for 250 steps each using first the steepest descent algorithm and second the conjugate gradient algorithm, prior to inserting the protein into the equilibrated bilayer patch. After conversion to all-atom representations, the lipids, waters, and ions were energy minimized for 100 steps using the conjugate gradient algorithm. To ensure that no lipids overlapped with ring residues of the protein, dummy atoms were placed in the center of the rings during a second 100-step energy-minimization. Subsequently the system was energy-minimized for 5000 steps using the conjugate gradient algorithm with positional restraints on the protein backbone, the non-hydrogen atoms of the side chains, and the O atoms of cavity waters. During a final 500 steps of conjugate gradient minimization, we applied a protein center-of-mass restraint and dihedral restraints on the φ and ψ and χ_1_ angles, with a force constant of 256 kJ/mol. The ions in the protein binding sites and the Arg104–Glu493 salt bridge were restrained to a minimum number of coordinating interactions using the COLVAR module (46). Specifically, ion coordinating atom (Ct1 comprising Ala96 O, Asn101 O_δ1_, Ser336 O, Ser336 O_γ_, Asn368 O_δ1_; Ct2 comprising Gly94 O, Val97 O, Leu434 O, Ser438 O_γ_; and Cl^−^ comprising Asn101 N_δ2_, Tyr121 O_ρι_, Gln332 N_ε2_, S336 O_γ_, S372 O_γ_) distances were set to ≤2.5 Å for Na^+^, ≤2.8 Å for K^+^, and ≤3.5 Å for Cl^−^, with a force constant of 10 kcal/mol and a minimum coordination number of 2. In the outward-occluded conformation, the Arg104–Glu493 salt bridge distance (between Arg104 N_ε_, Arg104 N_ρι1_, or Arg104 N_ρι2_ and E493 O_δ1_ or E493 O_δ2_) was restrained to ≤3.5 Å, with a force constant of 50 kcal/mol and a minimum coordination number of 1. All these restraints were maintained during six subsequent stages of molecular dynamics (MD) equilibration lasting >204 ns in total, while progressively reducing the force constants of the angle restraints to zero as follows: (1) φ/ψ force constant 256 kJ/mol and χ_1_ force constant 256 kJ/mol, (2) φ/ψ 256 kJ/mol and χ_1_ 64 kJ/mol, (3) φ/ψ 64 kJ/mol and χ_1_ 16 kJ/mol, (4) φ/ψ 16 kJ/mol and χ_1_ 4 kJ/mol, (5) φ/ψ 4 kJ/mol and χ_1_ 1 kJ/mol, and (6) φ/ψ 4 kJ/mol and no χ_1_ restraint. The salt bridge restraint was kept at a force constant of 50 kcal/mol during the first 4 ns, then progressively reduced by 10 kcal/mol per stage until reaching 10 kcal/mol at equilibration stage 6.

MD simulations were carried out using a 2 fs time step and trajectory output was saved every 4 ps for analyses. A temperature of 298 K was maintained using Langevin Dynamics, and pressure was kept at 1.01325 bar (oscillation time scale = 200 fs; damping time scale = 50 fs) using Nosé-Hoover Langevin piston pressure control (47). A cut-off distance of 12 Å with a switching function starting at 10 Å was used for van-der-Waals interactions. The particle-mesh Ewald method (48) was used for long-range electrostatic forces. Multiple production runs, each ∼0.5 µs long, were carried out without any restraints applied, but with different initial velocities. Final frames from each trajectory, as well as parameter and input files are provided as online supporting information (http://zenodo.org/).

### MD Simulation Analysis

Distance analyses were performed with the COLVARS module v2022-05-24 (46) in VMD v1.9.3 (49). Occupancy maps of ions and water were calculated in VMD using the volmap plugin v1.1.

### Whole-cell patch clamping

Whole-cell patch-clamp experiments were performed on HEK293 cells stably expressing SERT or transiently expressing SERT-437T and SERT-S438A, respectively. HEK293 cells, purchased from ATCC (# CRL-1573; ATCC, USA), were authenticated by STR profiling at the Medical University of Graz (Cell Culture Core Facility). These cells were then used to generate a stable line expressing SERT wildtype or they were transiently transfected with D437T or S438A, in serum-free DMEM using polyethyleneimine 25kDa (PEI) (Santa Cruz Biotechnology, Inc.; Santa Cruz, TX, USA) as the transfecting agent. The plasmid: PEI ratio was 1:3 (w/w). The cells were regularly tested for mycoplasma contamination by 4′,6-diamidino-2-phenylindole staining. These cells were grown in Dulbecco’s Modified Eagle Media (DMEM) supplemented with 10% heat-inactivated fetal calf serum (FBS), 0.6 µg ml^−1^ penicillin, 1 µg ml^−1^ streptomycin, and 100 μg ml^−1^ of geneticin/G418 for positive selection of transporter-expressing clones. None of the cell lines used in this study were included in the list of commonly misidentified cell lines maintained by the International Cell Line Authentication Committee.

24 hours prior to patching, the cells were seeded at a low density on PDL-coated 35 mm plates. Substrate-induced transporter currents were recorded under voltage clamp. Cells were continuously superfused with a physiological external solution that contained 163 mM NaCl, 2.5 mM CaCl_2_, 2 mM MgCl_2_, 20 mM glucose, and 10 mM 4-(2-hydroxyethyl)-1-piperazineethanesulfonic acid (HEPES) (pH adjusted to 7.4 with NaOH). A pipette solution mimicking the internal ionic composition of a cell contained 163 mM K-mesylate, 1 mM CaCl_2_, 0.7 mM MgCl_2_, 10 mM EGTA (ethylene glycol-bis(β-aminoethyl ether)-N,N,N’,N’-tetraacetic acid), and 10 mM HEPES (pH adjusted to 7.2 using KOH). In experiments in which we omitted intracellular potassium, we used the following solution: 163 mM NMDG-mesylate, 1 mM CaCl_2_, 0.7 mM MgCl_2_, 10 mM EGTA, and 10 mM HEPES (pH adjusted to 7.2 using KOH). For experiments in which we lowered the internal pH to 5.5, we used a potassium-free solution, where we replaced HEPES with MES (2-(N-morpholino)ethanesulfonic acid). The pH was adjusted to 5.5 using methanesulfonic acid (MsOH).

For patch clamp recordings, we employed the whole-cell configuration using an Axon 200B amplifier equipped with an Axon 1550 digitizer (both from Molecular Devices, LLC, San José, CA, USA). Recordings were sampled with 10 kHz. We quantified current amplitudes and associated kinetics using Clampfit software (Molecular Devices, LLC, San José, CA, USA). Passive holding currents were subtracted, and the traces were digitally filtered using an 80 Hz digital 8-pole Bessel low-pass filter.

5-HT or cocaine was applied using a perfusion system (Octaflow II, ALA Scientific Instruments, Inc., Farmingdale, NY, USA) that allowed for rapid and complete solution exchange around the cells within approximately 50 milliseconds. All experiments were conducted at room temperature (25°C) using a temperature control unit (cell microcontrol inline preheater, Green Leaf Scientific, Dublin, Ireland) connected to a temperature control system (PTC-20, npi electronic GmbH, Tamm, Germany). Both the patch-clamp holder and pipette electrode were operated via a PatchStar micromanipulator (Scientifica Ltd., Uckfield, East Sussex, United Kingdom).

## Acknowledgements

This research was supported by the Division of Intramural Research of the NIH, National Institute of Neurological Disorders and Stroke, NS003139 (to E.H. and L.R.F.), the National Natural Science Foundation of China (32071233 and 32371304 to YW.Z.), the Guangzhou City-University Joint Research Program (2023A03J0046 to YW.Z.), the Basic and Innovative Research Program for Guangzhou University Postgraduate Students (2022GDJC-M15 to Q.C.), the Austrian Science Fund/FWF (P36667 to W.S.) and the Vienna Science and Technology Fund/WWTF (LSC17-026 to M.F.). This work utilized the computational resources of the NIH High Performance Cluster, Biowulf. We thank Jessica Rana for researching possible computational strategies and José Faraldo-Gómez and Vanessa Leone for helpful discussions.

