## Supplemental Table and Figures for "Identification of the potassium binding site in serotonin transporter SERT"

**Table S1: Kinetic parameters for $APP^+$ uptake by SERT WT and Ct2-site mutants**

| | $K_m$ (µM) | $V_{max}$ (AFU) |
|---|---|---|
| WT | 2.12 ± 0.26 | 35.36 ± 1.33 |
| D437T | 1.81 ± 0.15 | 27.72 ± 2.08 |
| D437A | n.f. | n.f. |
| S438A | 2.01 ± 0.14 | 34.83 ± 1.19 |

Kinetic analysis for $APP^+$ uptake was performed with HEK 293T cells stably expressing SERT WT or mutant by incubating with $APP^+$ over a concentration range of 0.1–10 µM in KRH buffer containing 20 mM HEPES, pH 7.4, 120 mM NaCl, 1.3 mM KCl, 2.2 mM $CaCl_2$, 1.2 mM $MgSO_4$, and 0.1% (w/v) glucose for 5 min at 22°C. $APP^+$ accumulation in the cells was measured (n = 3). n.f., nonfunctional.

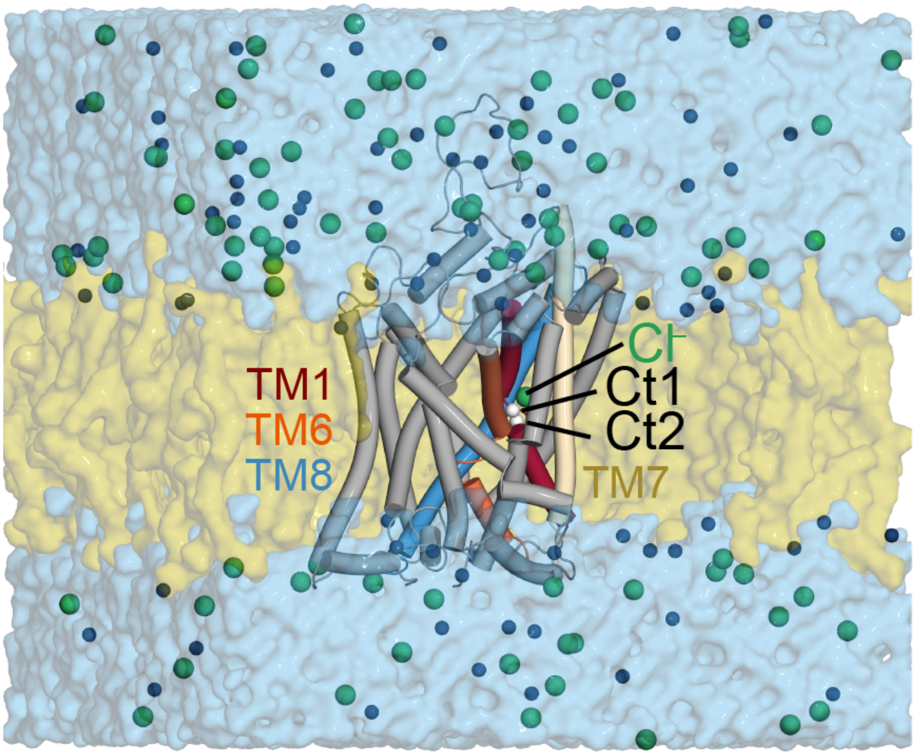


**Figure S1**. **SERT simulation system and binding site locations**. The simulation systems comprise a water solution (blue surface) at physiological salt concentration ($Na^+$ as dark blue spheres, $Cl^-$ as green spheres) and a homogenous POPC lipid bilayer (olive brown surface) in which outward-occluded SERT (cartoon helices) is embedded. The two cation sites being investigated, named Ct1 and Ct2, are shown as white spheres in the core of the transmembrane region, surrounded by TM1 (red), TM8 (blue), TM6 (orange) and TM7 (wheat). The remainder of the protein is colored gray.

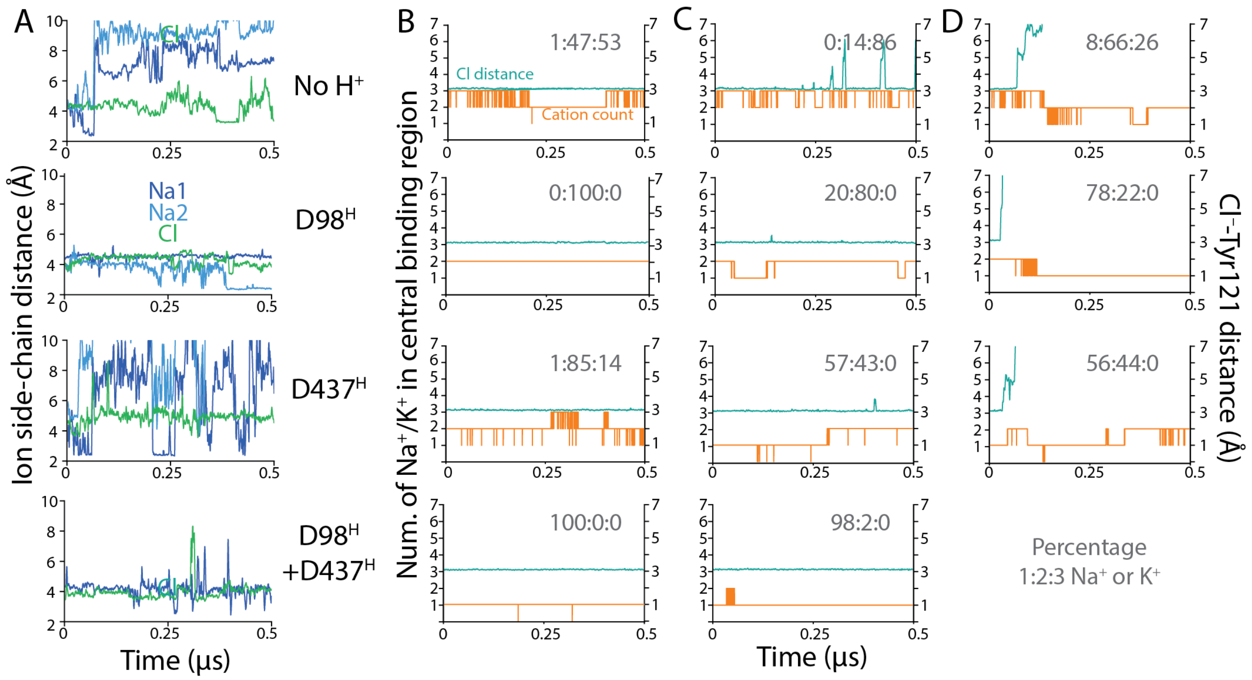


**Figure S2: Interactions of ions with SERT are stochastic when the intracellular pathway is open**. During MD simulations of the inward-facing conformation of SERT (n = 1 to 4 for each configuration), Asp98 and Asp437 were either both charged (top row), only Asp98 was protonated (second row), only Asp437 was protonated (third row), or both were protonated (bottom row). (**A**) **Stochastic unbinding can occur from both cation sites, as well as from the $Cl^-$ site**. Ion distances to example binding site residues are plotted as a function of simulation time for representative simulations. Distances were measured to the side-chain O of Asn101 (to a $Na^+$ placed in Ct1, dark blue), the closest of the two side-chain O atoms of Asp437 (to the $Na^+$ placed in Ct2, light blue), and the side chain N of Asn368 (to the $Cl^-$ ion, green). (**B-D**) **The central pocket holds 3 cations (including covalently-linked protons at Asp98 and/or Asp437), when $Cl^-$ is bound**. After cation unbinding events, the net charge is balanced by stochastic binding of $Na^+$ or $K^+$ ions from the cytoplasmic solution. The total number of $Na^+$ or $K^+$ ions occupying the S1 region is plotted as a function of simulation time (orange). The S1 region is defined as a radius of 10 Å using the center of mass of the Cα atoms of Asp98 and Asp437 as a reference point. Ions were counted if they were initially bound, or if they approached within 6 Å of the same reference point during the trajectory. Distances of the bound $Cl^-$ ion to the side chain Oγ of Tyr121 are shown for reference (green lines). Columns (B) and (C) provide examples of $Cl^-$ remaining bound for simulations with either 150 mM NaCl (B), or 130 mM KCl and 10 mM NaCl (C). Column (D) illustrates $Cl^-$ ion unbinding events, upon which the net cation count reduces to ~2. For each example, the percentage of the trajectory in which 1, 2, or 3 $Na^+$/$K^+$ ions occupy the pocket is provided (gray text).

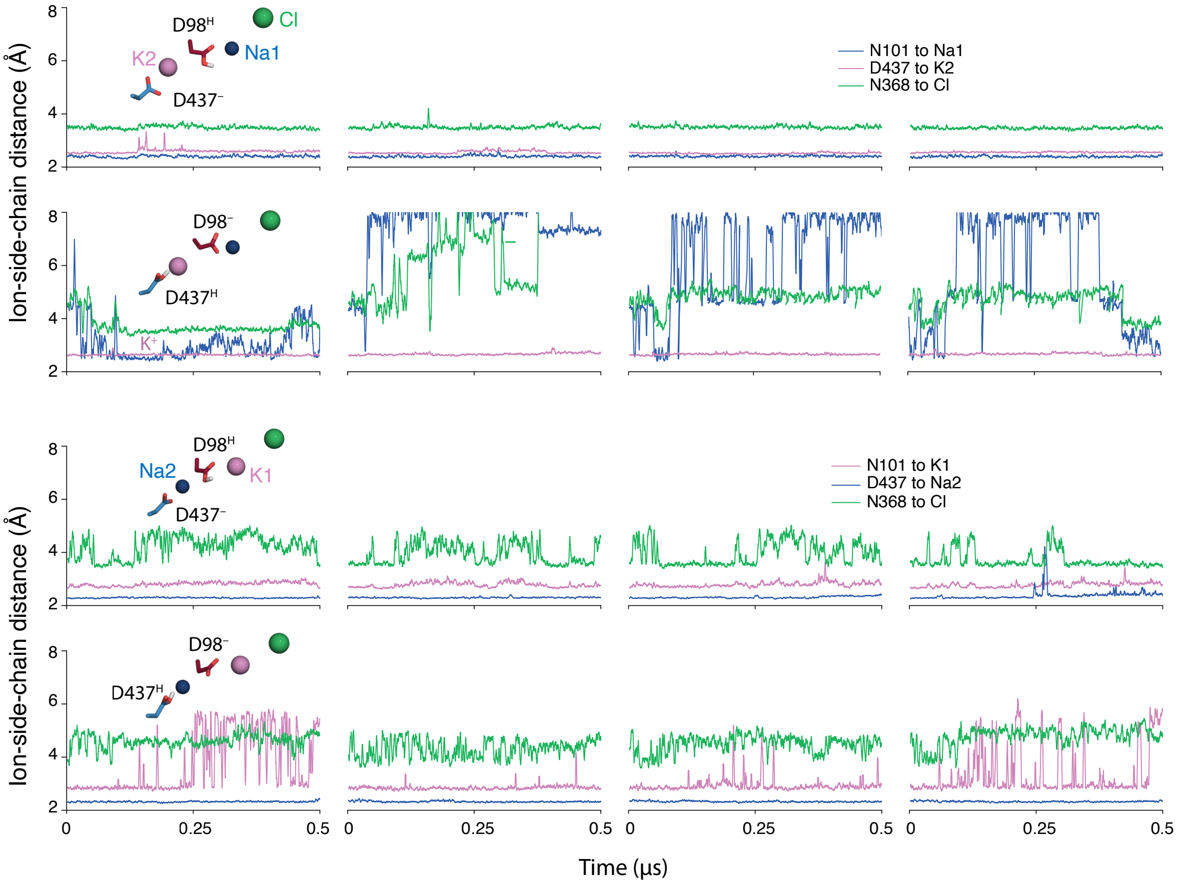


**Figure S3. Ion distances to selected binding site residues as a function of simulation time for the occluded conformation of SERT.** Each row shows the time courses of n = 4 simulation replicas starting with a conformation in which both pathways are closed. Inset structures are representative snapshots from the respective simulation. The configuration in which D98 is protonated, D437 is charged, with $Na^+$ (blue) in Ct1 and $K^+$ (violet) in Ct2 (first row) is predicted to have the most stable ion coordination overall. Large fluctuations occur when D98 is charged (rows 2 and 4) and when $K^+$ is bound to Ct1 (rows 3 and 4). Distances are measured to the side-chain O of Asn101 (to the ion initially placed in Ct1), the minimum to the two side-chain O atoms of Asp437 (to the ion initially placed in Ct2), or the side chain nitrogen of Asn368 (to the $Cl^-$ ion).

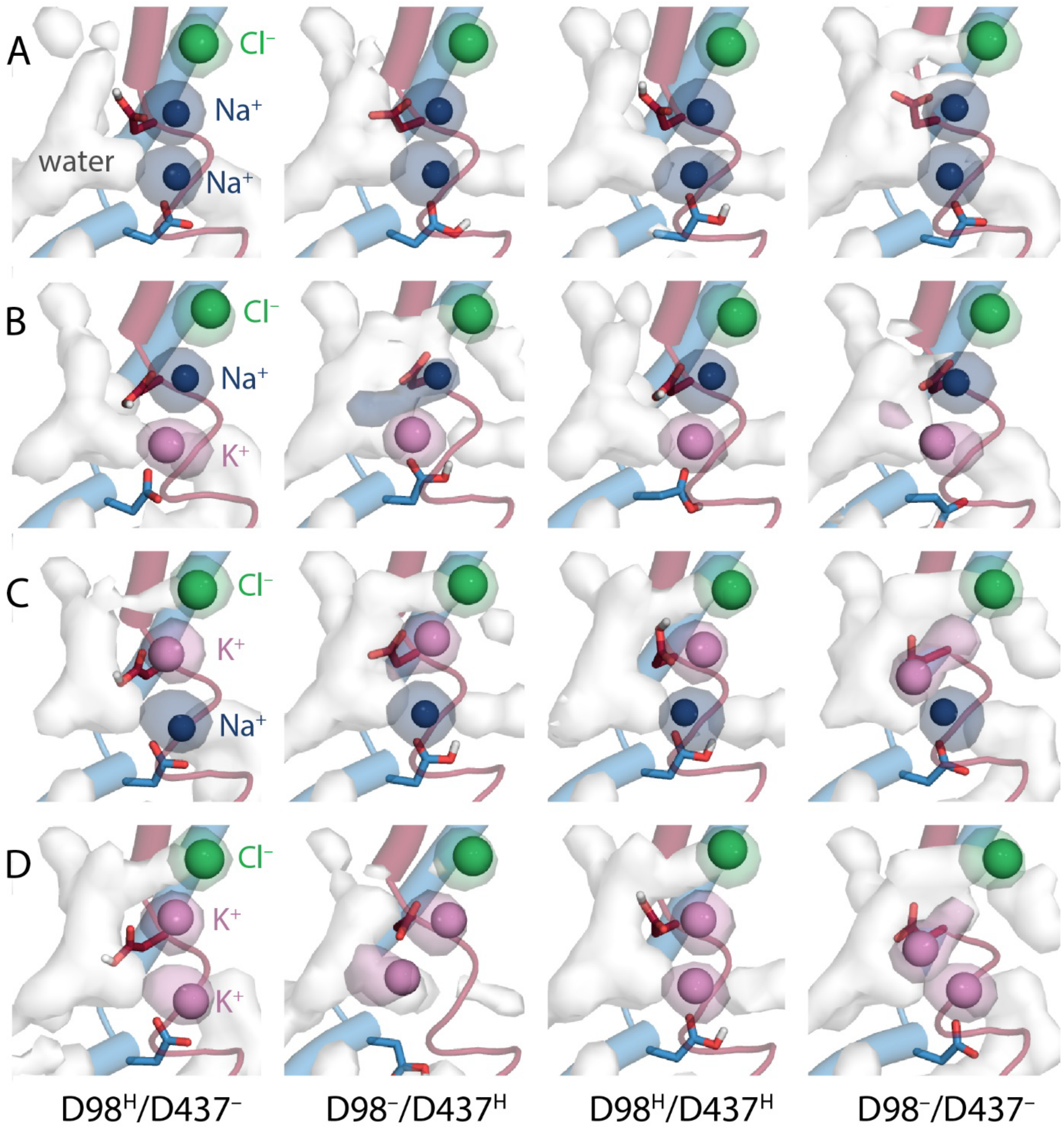


**Figure S4. Ion and water occupancy during molecular dynamics simulations of SERT in the absence of 5-HT**. Densities are shown for simulations in which cation site 1 (Ct1) was occupied with either $Na^+$ (A, B) or $K^+$ (C, D), and where cation site 2 (Ct2) was occupied with $Na^+$ (A, C) or $K^+$ (B, D). The protonation states of Asp98 and Asp437 were also varied, with either a single proton (left and center left columns), no protons (right-hand column) or two protons (center right column). Occupancy maps (surfaces) were computed over repeated simulation trajectories (n = 4), and are shown for water in white, $Na^+$ in dark blue, $K^+$ in violet, and $Cl^-$ in green. The four panels shown in Figure 3C are included here for completeness.

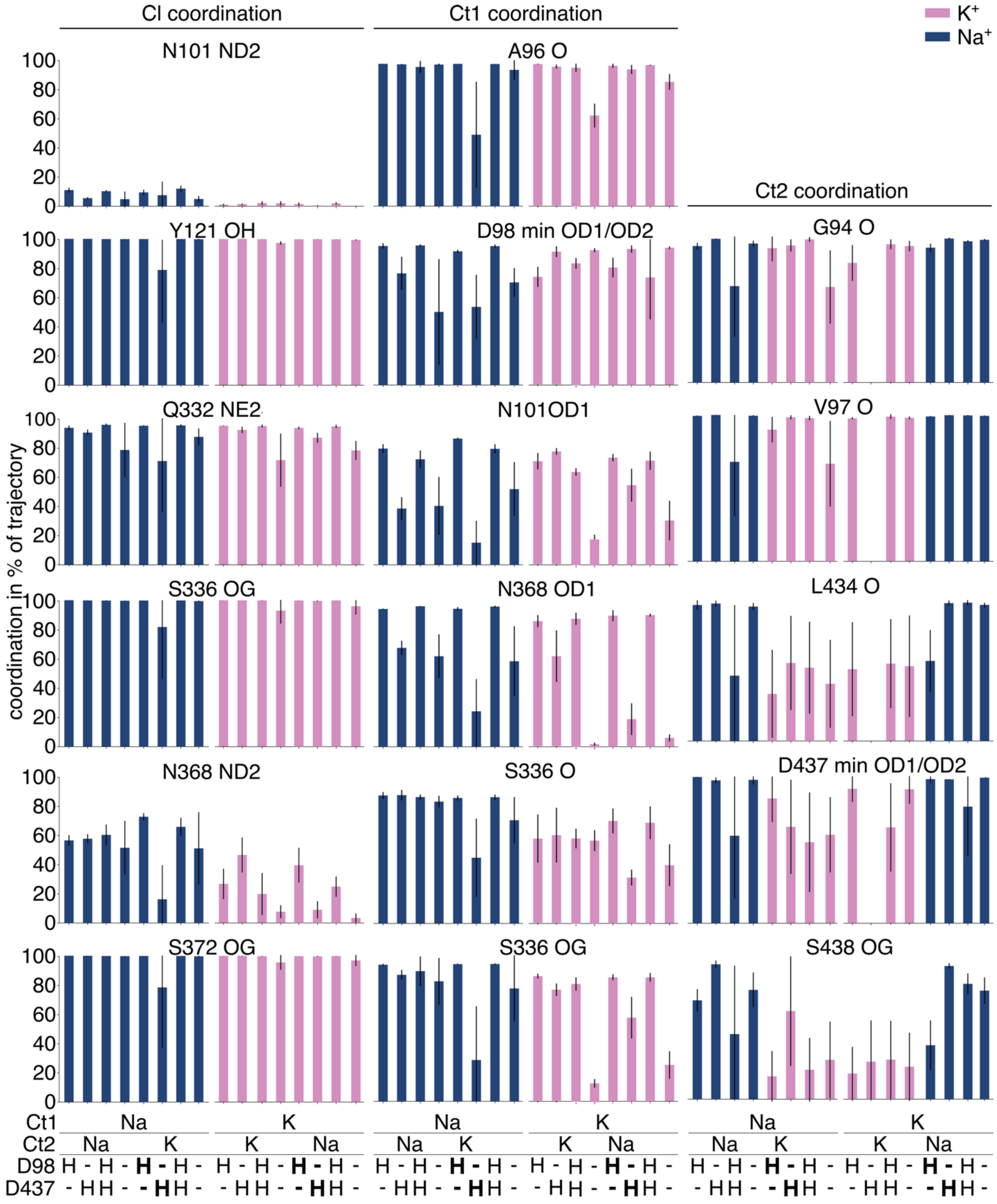


**Figure S5. Coordinating distances for all simulation systems.** Values are mean (and standard deviation) of the percentage of the simulation time that the ion ($Cl^-$, Ct1, or Ct2, from left to right), colored blue and purple for $Na^+$ and $K^+$, respectively, is within coordination distance of the indicated atom. All combinations of bound cations and protonation states for Asp98 and Asp437 were tested. See legend to Figure 3 for more details.

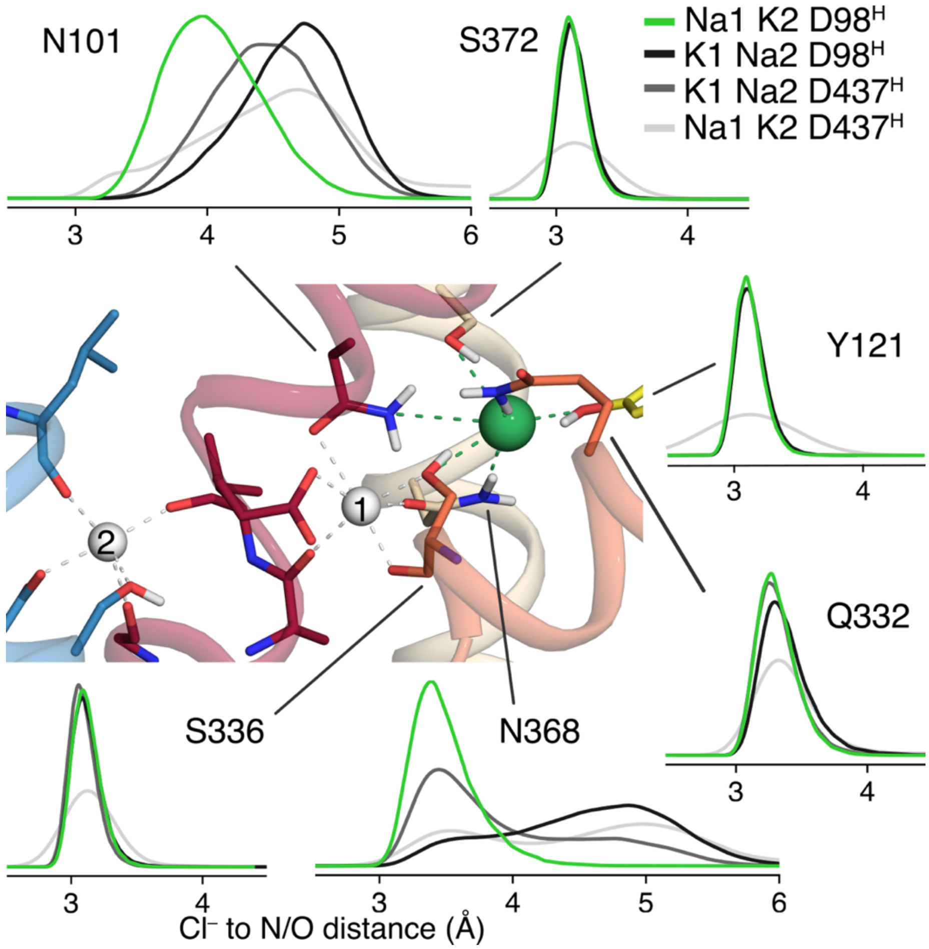


**Figure S6: Analysis of Cl⁻-protein interactions during molecular dynamics simulations of SERT with different cations at the Ct1 and Ct2 sites, and different protonation states for Asp98 and Asp437**. Six interactions, shown as dashed green lines in the central structural snapshot, were tracked using the distances from side-chain O or N atoms to the Cl⁻ ion. Distributions are averaged over multiple repeat trajectories. Interactions were stable for all coordinating residues when the Ct1 site was occupied with a $Na^+$ ion, the Ct2 site contained a $K^+$ ion, and Asp98 was protonated (green). Interactions with Asn101 and Asn386 were disrupted by a $K^+$ ion at Ct1 (black; dark gray), or by deprotonating Asp98 (light gray). The formatting for the central figure follows that in Figure Y. These figures indicate that the molecular origin of the preference of $K^+$ for Ct2 is destabilization of the protein-Cl⁻ ion interactions.

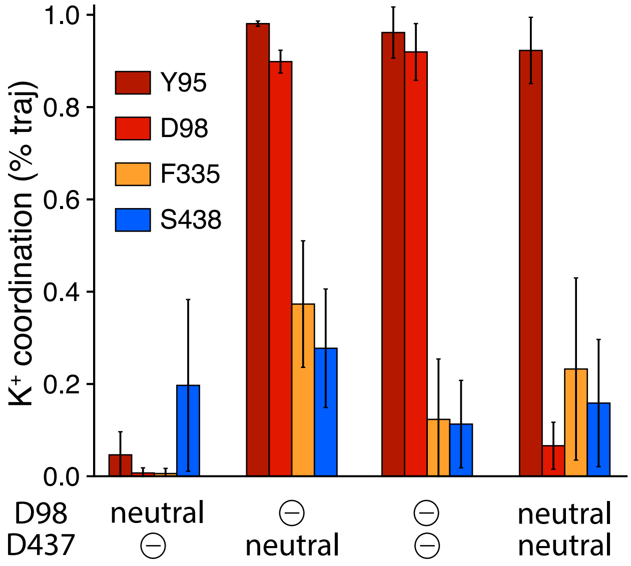


**Figure S7: Interactions formed by a $K^+$ ion placed in the central substrate binding site of SERT with different protonation states of Asp98 and Asp437**. Coordination was assessed as the percentage of frames over all trajectories that the ion was within a cutoff distance from the coordinating atoms. Error bars reflect the standard deviation computed over repeated trajectories (n = 10 for simulations with a single added proton, and n = 6 for doubly charged or doubly protonated versions). Distances were measured to the center of mass of the aromatic ring of Tyr95 in TM1; the minimum of the two side-chain carboxyl O atoms of Asp98; the backbone carbonyl O atom of Phe335 in TM6; or the hydroxyl O atom of Ser438 in TM8 with thresholds of <6 Å for aromatic groups and <2.9 Å for O atoms.

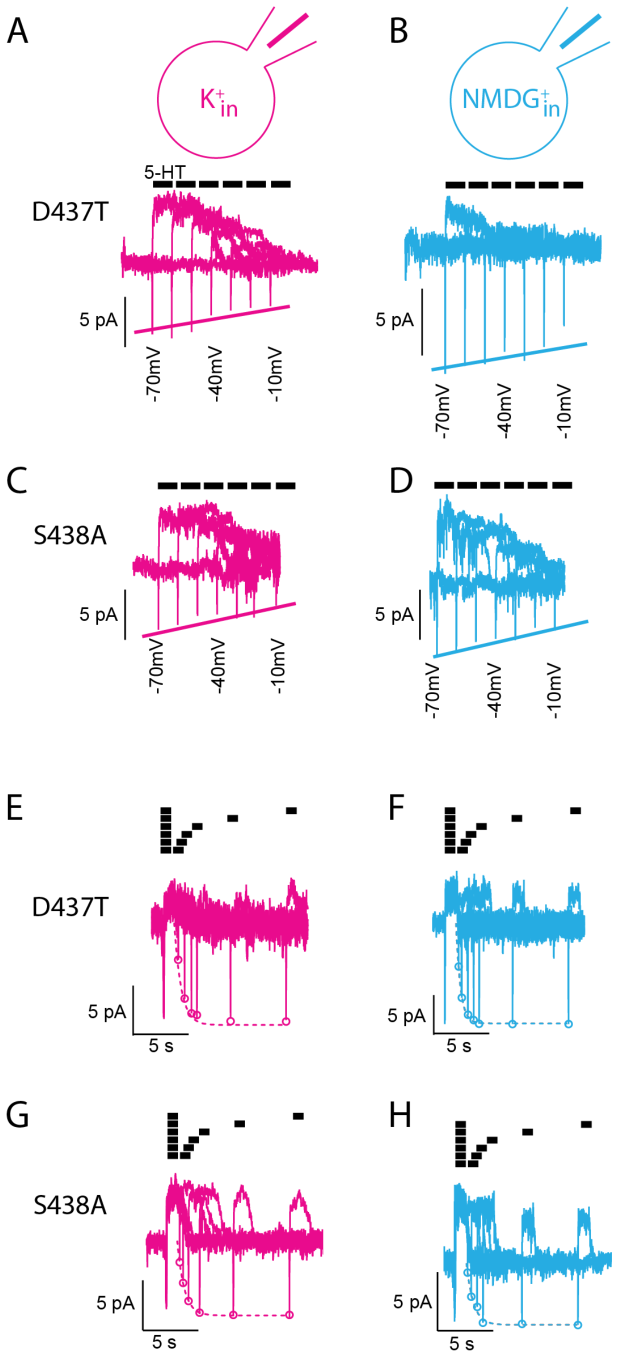


**Figure S8: The substrate-induced current through D437T and S438A is insensitive to $K^+_{in}$.** (**A-D**) Representative traces of peak currents obtained from SERT mutants D437T (A, B) and S438A (C, D) in the presence of 160 mM $K^+_{in}$ (A, C) or 160 mM $NDMG^+_{in}$ (B, D) recorded at voltages ranging from –60 mV to +10 mV. Solid lines reflect the fits shown in Figure 5D for D437T and Figure 5E for S438A. (**E-H**) Representative current traces resulting from a two-pulse protocol recorded from SERT mutants D437T (E, F) and S438A (G, H). The intracellular solution contained 160 mM $K^+$ (E, G) or 160 mM $NMDG^+$ (F, H). Dashed lines and circles correspond to the data in Figure 5I for D437T and Figure 5J for S438A.

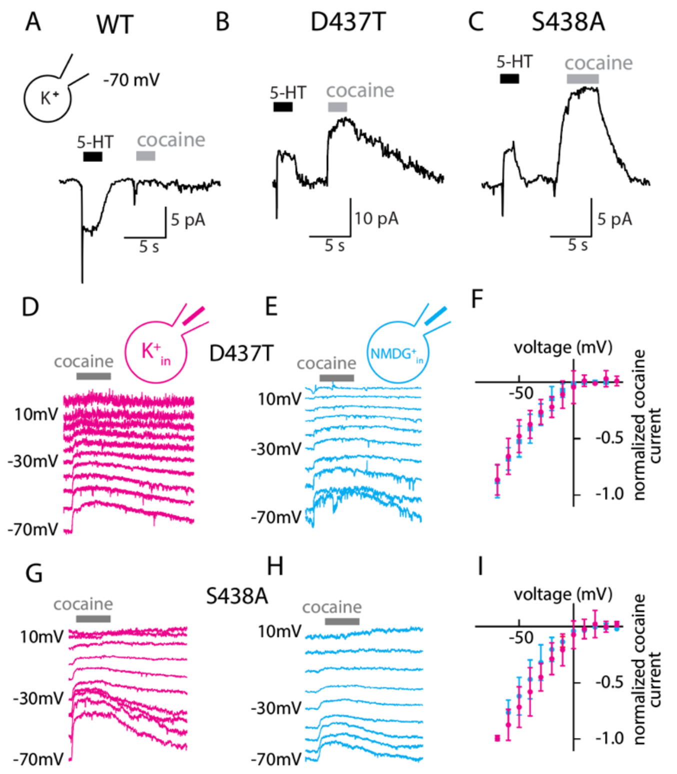


**Figure S9: Ct2 site mutants mediate a leak current that is insensitive to K$^+_{in}$.** (**A**) Representative current induced by 30 µM 5-HT recorded from wild type SERT at –70 mV in the presence of 160 mM K$^+_{in}$. The subsequent application of a saturating concentration of cocaine (i.e., 30 µM) gave rise to a small peak current but did not provide evidence for the existence of a substrate-independent leak current carried by wild type SERT. (**B, C**) Representative currents recorded at –70 mV from D437T (B) and S438A (C) in the presence of 160 mM K$^+_{in}$. Application of 30 µM 5-HT induced a peak current, which was followed by an outward directed steady current. After removal of 5-HT from the bath solution, the currents of the two mutants relaxed to baseline levels. Subsequent administration of 30 µM cocaine, an inhibitor that stabilizes the outward-open state and is expected to inhibit charge flux, revealed that this mutant carries a substrate-independent leak current. Thus, the outward-directed current seen in the presence of 5-HT (B, C) likely results from a substrate-induced (partial) block of a constitutive leak current through these mutants. A possible origin of this behavior is that the mutants have developed the ability to transport Na$^+$ in the absence of substrate, or rather, have lost the tight coupling observed in the wild type protein. (**D, E**) Representative traces of leak currents through D437T in the presence of 160 mM K$^+_{in}$ (E) or 160 mM NMDG$^+_{in}$ (F) recorded in the voltage range between –70 mV and +10 mV. The leak current was defined by the application of 30 µM cocaine. (**F**) Normalized current amplitudes measured from D437T as a function of the membrane potential in the presence of 160 mM K$^+_{in}$ (n = 13, circles) and NMDG$^+_{in}$ (n = 7, squares), respectively. (**G, H**) Representative traces of leak currents through SERT S438A in the presence of 160 mM K$^+_{in}$ (H) or 160 mM NMDG$^+$ (I) recorded in the voltage range between –70 mV and +10 mV. (**I**) Normalized current amplitudes measured from S438A as a function of the membrane potential in the presence of 160 mM K$^+_{in}$ (n = 14, circles) and NMDG$^+_{in}$ (n = 9, squares), respectively. The amplitude of the leak current decreases at more positive potentials for both mutants. Moreover, the current-voltage relation was the same in the presence and absence of 160 mM K$^+_{in}$ for both D437T (F) and S438A (I), providing support for the idea that neither mutant can bind K$^+$.

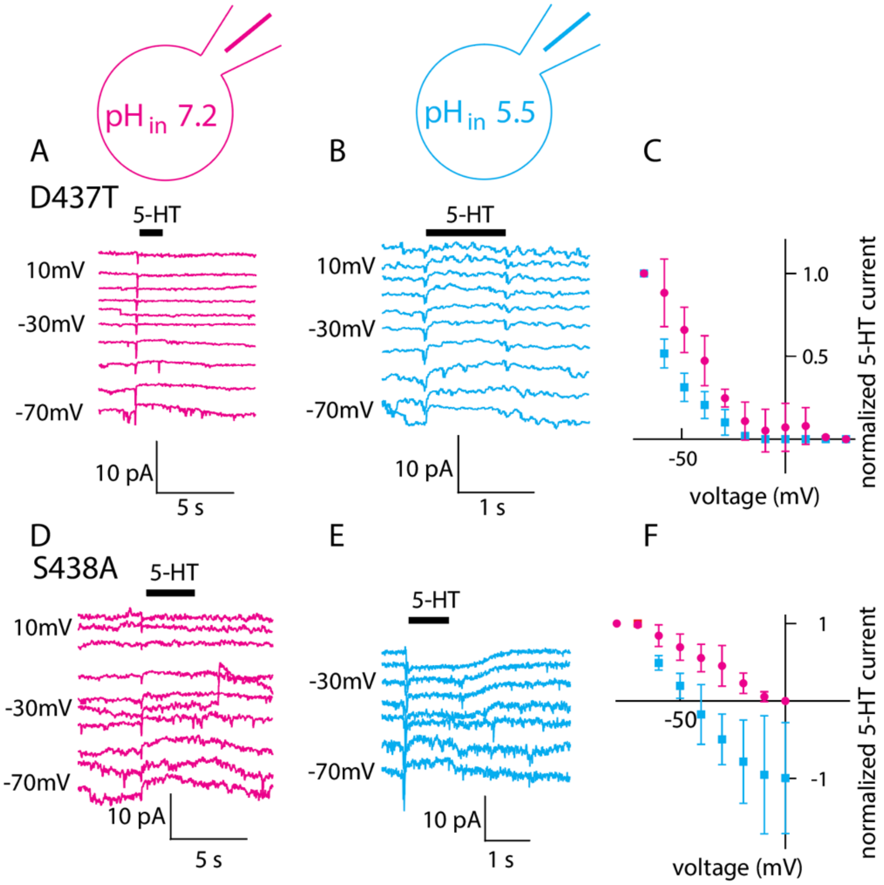


**Figure S10. The proton dependence of SERT requires Asp437**. (**A, B**) Representative traces of currents induced by 30 µM 5-HT recorded from D437T in the voltage range from –70 mV to 10 mV. In this experiment, the intracellular solution contained 160 mM $NMDG^+$, and its pH was set to 7.2 (A) or 5.5 (B). At the most negative potentials, the peak currents were followed by outward-directed steady currents, consistent with leak currents partially blocked by the substrate (cf. Fig. S9). (**C**) Normalized amplitudes of the 5-HT-induced current of D437T plotted as a function of membrane voltage with the intracellular pH adjusted to 7.2 (n = 6, circles) and 5.5 (n = 5, squares). Increasing the intracellular $H^+$ concentration had minimal impact on the current-voltage relationship. Moreover, at both tested pH values, the substrate-induced steady currents were all positive, i.e., outward directed, and did not reverse direction at less negative applied voltages, indicating that the removal of the acidic side chain modifies the response of SERT to low $pH_{in}$. (**D, E**) Representative traces of currents induced by 30 µM 5-HT recorded from S438A in the voltage range from –70 mV to 10 mV. The intracellular pH in this experiment was 7.2 (D) or 5.5 (E). In the presence of $pH_{in}$ 7.2, the currents were similar to the currents through D437T. Unlike D437T, the currents elicited by S438A were sensitive to the intracellular pH. (**F**) Normalized amplitudes of the 5-HT-induced current measured from S438A as a function of voltage at an intracellular pH of 7.2 (n = 9, circles) or 5.5 (n = 9, squares). At $pH_{in}$ 5.5, the current was outwardly directed at –70 mV and reversed direction at around –45 mV. Thus, the voltage dependence manifests as a higher apparent $H^+$ affinity at positive potentials, following the same trend as previous measurements of the peak currents elicited by wild type SERT.

**Legend to Movie S1: A $K^+$ ion reproducibly binds to the Ct2 site in the occluded conformation of SERT.** Molecular dynamics simulation trajectory of SERT with Asp98 protonated and Asp437 charged. In this condition, a $K^+$ ion reproducibly binds to the Ct2 site in the occluded conformation. Four key transmembrane helices of SERT are shown as cylinders (TM1 in red, TM6 in orange, TM7 in wheat, and TM8 in cyan), with key residues shown as sticks (see Figure 1B). The bound $Cl^-$ (green), $Na^+$ (blue) and K+ (purple) ions are shown as spheres. The simulation was initiated with the $K^+$ ion in the central S1 substrate binding site close to Asp98, which was protonated. See Fig. 4 for reproducible binding traces. Each trajectory is 0.5 μs long.

**Legend to Movie S2: Binding of $K^+$ at the Ct2 site of SERT is dependent on the protonation state of Asp98.** Molecular dynamics simulation trajectory of SERT with Asp98 deprotonated and Asp437 protonated. The simulation was initiated with the $K^+$ ion in the central S1 substrate binding site close to Asp98, which was charged. See description to Move S1 for details, and Fig. 4 for reproducible binding traces. Each trajectory is 0.5 μs long.
